# Genome-wide single cell annotation of the human protein-coding genes

**DOI:** 10.1101/2022.08.03.502627

**Authors:** Max Karlsson, María Bueno Álvez, Mengnan Shi, Loren Méar, Rutger Schutten, Feria Hikmet, Andreas Digre, Borbala Katona, Jimmy Vuu, Martina Bosic, Evelina Sjöstedt, Fredrik Edfors, Per Oksvold, Kalle von Feilitzen, Martin Zwahlen, Mattias Forsberg, Fredric Johansson, Jan Mulder, Tomas Hökfelt, Younglun Luo, Lynn Butler, Wen Zhong, Adil Mardinoglu, Åsa Sivertsson, Fredrik Ponten, Cheng Zhang, Cecilia Lindskog, Linn Fagerberg, Mathias Uhlén

## Abstract

An important quest for the life science community is to deliver a complete annotation of the human building-blocks of life, the genes and the proteins. Here, we report on a genome-wide effort to annotate all protein-coding genes based on single cell transcriptomics data representing all major tissues and organs in the human body, integrated with data from bulk transcriptomics and antibody-based tissue profiling. Altogether, 25 tissues have been analyzed with single cell transcriptomics resulting in genome-wide expression in 444 single cell types using a strategy involving pooling data from individual cells to obtain genome-wide expression profiles of individual cell type. We introduce a new genome-wide classification tool based on clustering of similar expression profiles across single cell types, which can be visualized using dimensional reduction maps (UMAP). The clustering classification is integrated with a new “tau” score classification for all protein-coding genes, resulting in a measure of single cell specificity across all cell types for all individual genes. The analysis has allowed us to annotate all human protein-coding genes with regards to function and spatial distribution across individual cell types across all major tissues and organs in the human body. A new version of the open access Human Protein Atlas (www.proteinatlas.org) has been launched to enable researchers to explore the new genome-wide annotation on an individual gene level.

## Introduction

The Human Genome Project has profoundly transformed and accelerated our understanding of human biology and its implications for disease. In the two papers from 2001 (*1, 2*) the completion of the genome sequence was described, but it is noteworthy that 15% of the genome was missing due to technical limitations. A multitude of efforts have since focused on filling these gaps, but the reference genome published in 2013 (*3*) still lacks 8% of the complete sequence. Recently, about 200 million bases of the haploid genome was added (*4*) using a new technology platform allowing sequencing of up to 20,000 bases, resulting in a nearly completed assembly of the human genome. The new update has given the research community a valuable blueprint of the hereditary genetic information in humans, including its variability across various populations (*5, 6*).

For a deeper understanding of human biology and disease, it is however necessary to annotate the function and spatial distribution of the primary effectors in biology; the genes and the proteins. The genes provide a “parts list” of the corresponding proteins necessary to make up humans and the proteins provide the functional units of human life. An important aim is thus to gain a holistic understanding of all proteins encoded by the genome on an individual gene level, but also to substantially increase our understanding of their structure, function, interactions and spatial distribution. A large number of initiatives has been launched aimed to provide a comprehensive view of all proteins and expression of the corresponding protein-coding genes with the objective to generate maps and annotate the spatial distribution and function of all proteins encoded by the human genome.

The Human Cell Atlas (*7*) uses single cell transcriptomics to study the gene expression profiles on the RNA level across single cells from tissues and organs. Similarly, the Tabula Sapiens consortium (*8*) has recently described the single cell levels in 24 organs across the human body and the Expression Atlas recently launched the Single Cell Expression Atlas for browsing and visualizing single cell data, which now contains data from more than 100 studies of human cells (*9*). These efforts have been complemented with more direct analysis of the proteins, including the OpenCell program (*10*) in which endogenous tagging is achieved using a brute-force CRISPR strategy to obtain a cartography of human cellular organization. In addition, the Human Proteoform Project (*11*) has been proposed to characterize the human protein variants (proteoforms), and the Human Proteome Project was launched (*12*) with a focus to create a more refined map of the human proteome using mass spectrometry. In parallel, several initiatives have been set up to take advantage of the many millions of antibodies towards human proteins now publicly available, pioneered by the Human Protein Atlas consortium (*13*) generating high-resolution protein localization data on a single cell level. More than 10 million bioimages are available in an open access resource (www.proteinatlas.org) showing the native protein location in various cells, tissues, and organs.

Here, we present a genome-wide annotation of the protein-coding genes based on single cell transcriptomics representing the major tissues and organs in the human body, integrated with bulk transcriptomics analysis of the corresponding tissues, and complemented with high-resolution antibody-based profiling to visualize single cells in intact tissue samples. Altogether, 25 tissues have been analyzed with single cell transcriptomics using a strategy described recently (*14*) involving pooling data from individual cell clusters to avoid issues with “drop-outs” related to genes with low expression. The analysis has allowed us to use a clustering approach to annotate all human protein-coding genes with regards to specificity and spatial distribution across single cell types corresponding to all major human tissues and organs. Grouping genes into local neighborhoods by dimensionality reduction and clustering has the potential to enhance the interpretability of gene expression data, find previously unknown relationships between genes, and highlight gaps in functional annotation for hypothesis generation. Each gene is thus classified into an expression cluster consisting of genes with similar single cell type expression across all the analyzed tissues. We here show that this approach can be used for genome-wide annotation of the human protein-coding genes integrating single cell transcriptomics with bulk transcriptomics and antibody-based profiling. A new version of the open access Single Cell Type section of the Human Protein Atlas is launched (www.proteinatlas.org/humanproteome/single+cell+type) to allow genome-wide exploration of individual genes and proteins with relation to spatial single cell distribution and function

## Results

### The single cell data included in the study

In this study, 25 tissues have been analyzed with single cell transcriptomics using datasets with the inclusion criteria as described previously (*14*) (**Supplementary data S1**-**S2**). Building on the previous analysis performed on scRNAseq data from 13 tissues; skin (*15*), eye (*16*), heart (*17*), small intestine (*18*), colon (*19*), rectum (*18*), kidney (*20*), liver (*21*), pancreas (*22*), lung (*23*), placenta (*24*), testis (*25*), and prostate (*26*), we here added 12 additional tissues; brain (*27*), skeletal muscle (*28*), esophagus (*29*), stomach (*29*), adipose tissue (*30*), bronchus (*31*), lymph node (*29*), bone marrow (*29*), spleen (*29*), breast (*32*), endometrium (*33*) and ovary (*34*), as well as peripheral blood mononuclear cells (PBMCs) (*35*), summarized in **Fig. 1A**. The additional tissues cover organ systems and unique single cell types not represented in the previous version, such as adipocytes, brain neuronal and glial cells, skeletal myocytes, gastric glands as well as specialized cells in the female reproductive system (mammary gland, endometrium, and ovary). Altogether, the dataset covers all major tissues and organs in the human body, and the combined data represents single cell analysis from a total of 566,108 individual cells.

**Figure 1.**
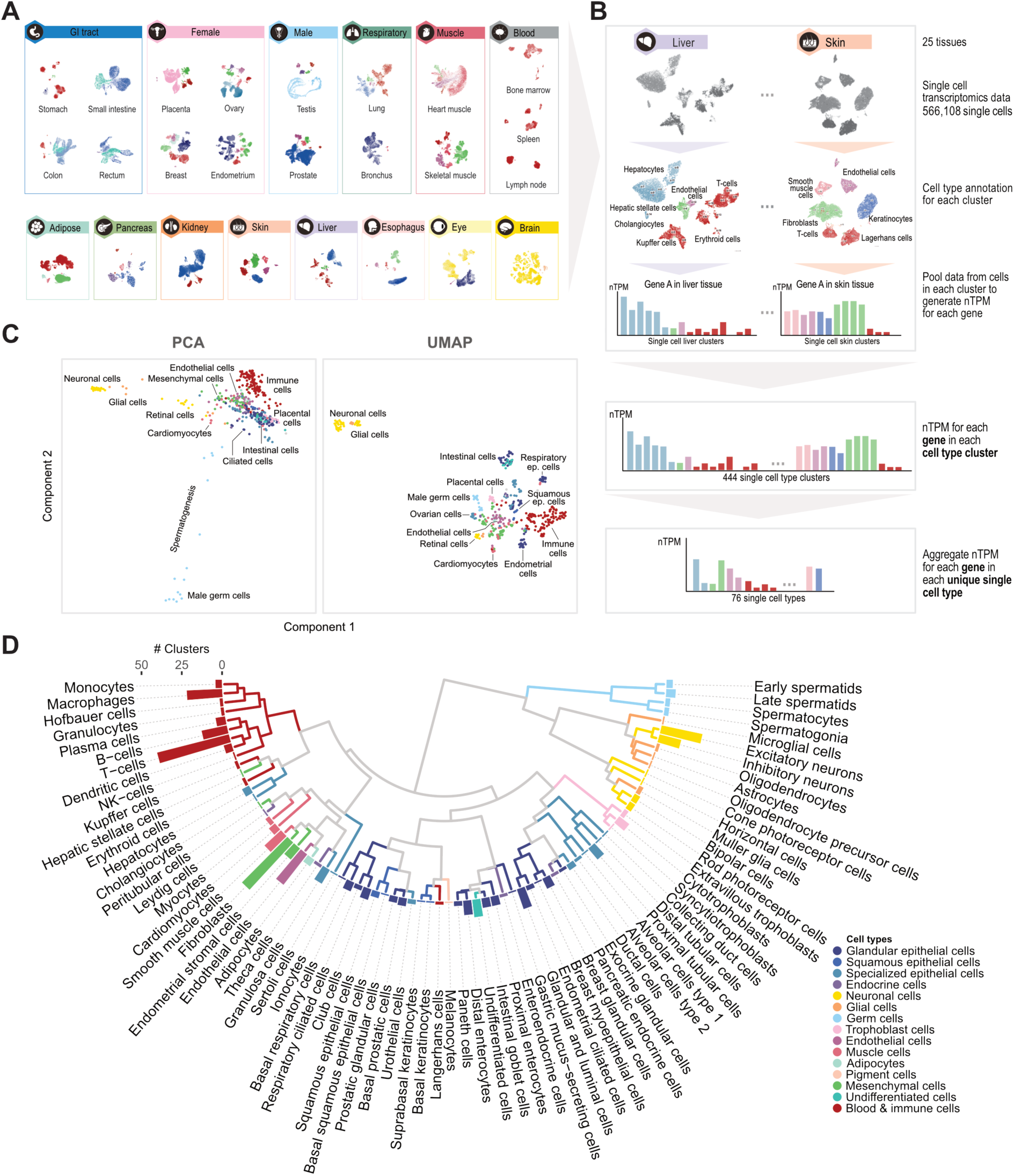
Overview of the single cell transcriptomics analysis. (**A**) Single cell transcriptomics datasets from 25 human tissues collected for the study. Cells from each dataset are displayed in a UMAP and colored by cell identity. (**B**) Pooling strategy used to calculate normalized transcripts per million (nTPM) expression values for each gene in each unique single cell type. (**C**) Principal component analysis (PCA) (left) and UMAP (right) visualization based on expression data for the 444 single cell type clusters. (**D**) Dendrogram of the 76 unique single cell types based on hierarchical clustering using Spearman correlation. Dendrogram branch lengths are scaled to emphasize connections, and are colored if all down-stream cells belong to the same cell type group. Bars indicate the number of cell type clusters pooled for each of the unique single cell types.

### The pooling strategy for analysis of single cell data

The scheme outlined in **Fig. 1B** was followed for all of the single cell datasets. For each of the 25 tissues, the raw data was quality controlled and cells with similar expression were grouped into clusters with each cluster representing a specific cell type. Between eight (lymph node) and 45 (brain) clusters were identified for each tissue to yield altogether 444 individual clusters which were manually annotated based on expression of established cell type marker genes as well as correlation with single cell type protein expression pattern based on immunohistochemistry. The data from each cluster was pooled to generate normalized transcript per million (nTPM) for each gene as described before (*14*). By combining the results from all 25 tissues, the expression profile for each gene across the 444 single cell type clusters was calculated. Since many of the single cell types were found across several tissues, such as immune cells, neurons, fibroblasts and endothelial cells, the 444 single cell types were aggregated into 76 consensus single cell types. In the open-access Single Cell Type section of the Human Protein Atlas (www.proteinatlas.org), the expression profiles (nTPM) for all genes are displayed across the individual 444 single cell type clusters and the 76 consensus single cell types.

### The transcriptional landscape of single cell types

The overall similarity in expression profiles of cell types were visualized by dimensional reduction analysis using Principal Component Analysis (PCA) or Uniform Manifold Approximation and Projection (UMAP). The results (**Fig. 1C**) for all the 444 identified single cell type clusters show that according to PCA, spermatids in testis and neuronal cells are the most distant cell types, while neuronal cells are most distant according to UMAP, implying that these cell types have the most unique expression profiles as compared to all the other cell types. For most single cell types, expected results were found, as for example the close similarity of expression for the various immune cells. The resulting expression profiles for each of the 76 consensus single cell types were analyzed using hierarchical clustering visualized in a dendrogram (**Fig. 1D** **and Fig. S1A)** also showing the number of tissues containing a specific cell type aggregated into the consensus cell types. Each of the 76 consensus single cell types were categorized into 15 major groups with related expression profiles, as exemplified by immune cells, epithelial cells, neuronal cells and endocrine cells, respectively.

### Bulk transcriptomics analysis of human tissues

In addition to single cell transcriptomics analysis based on single cell RNA sequencing (scRNA-seq), this study also includes bulk transcriptomics analysis of human tissues based on RNA-seq corresponding to 54 different tissue types aggregated into a consensus dataset of 36 tissues (**Supplementary data S3**). The bulk transcriptomics analysis covers all 25 tissues used for the single cell transcriptomics analysis. The consensus bulk transcriptomics dataset is normalized based on raw data that was either generated “in-house” as described earlier (*13, 36, 37*) or by the GTEx consortium (*5, 38*). The “in-house” data was complemented with an extensive new bulk transcriptomics dataset based on microdissection of human brain regions, totaling 1324 brain samples covering 206 subregions of the brain collected using the punch needle approach (*39*). A PCA analysis based on all 11,948 tissue samples shows that samples from testis, brain and muscle have the most distinct expression profiles, and a UMAP analysis further shows that samples from the same tissues are tightly grouped and separated from other tissue types, with similar or related tissues being in closer proximity (e.g. adipose and breast tissue) (**Fig. S2**). The relationship between different tissues based on bulk transcriptomics show similar clustering (**Fig. S2**) as compared to the single cell transcriptomics analysis (**Fig. 1C-D**) with testis and brain as outliers.

### The human secretome and membrane proteome

An important analysis for annotation of the human proteome is to determine the fraction of the total expression in each single cell type devoted to proteins with different localization in the cell, such as secreted, intracellular or membrane-bound. The results in **Fig. 2A** covering the 76 single cell types based on single cell transcriptomics show that for most of the cells, the majority (62-75%) of the gene expression corresponds to genes predicted to encode intracellular proteins, while less expression corresponds to genes encoding secreted (4-10%) or membrane-bound (18-26%) proteins. In this figure, the cell types are ordered according to the result of hierarchical clustering based on the Spearman distance between their expression profiles. Cell types with known functions associated with secretion, have as expected a much larger fraction of the transcriptome encoding secreted proteins. For example, plasma cells, the immune cells responsible for producing and secreting immunoglobulins, commit 55% of their total transcripts to genes with at least one secreted gene product; while for liver hepatocytes, which secrete large amounts of proteins to blood, such as the main plasma constituent albumin, the corresponding number is 39%. Another example is pancreatic exocrine glandular cells, with 35% of their transcripts corresponding to genes with secreted gene products, out of which approximately half (15%) relates to only three proteins secreted in large amounts; the pancreatic stone protein or Lithostathine-1-alpha, presumed to inhibit calcium carbonate precipitation (*40*) and the trypsin proteases PRSS2 and PRSS1 that act as digestive enzymes. High expression of all three proteins in pancreatic exocrine glandular cells is supported by the antibody-based tissue profiling (**Fig. S3**). The alveolar cells type 2 (AT2), which produce the pulmonary surfactant proteins crucial for the air-liquid interface, commit 26% of the expression to genes with secreted isoforms, out of which half (13%) corresponds to a single protein, the Surfactant Protein C (SFTPC).

**Figure 2.**
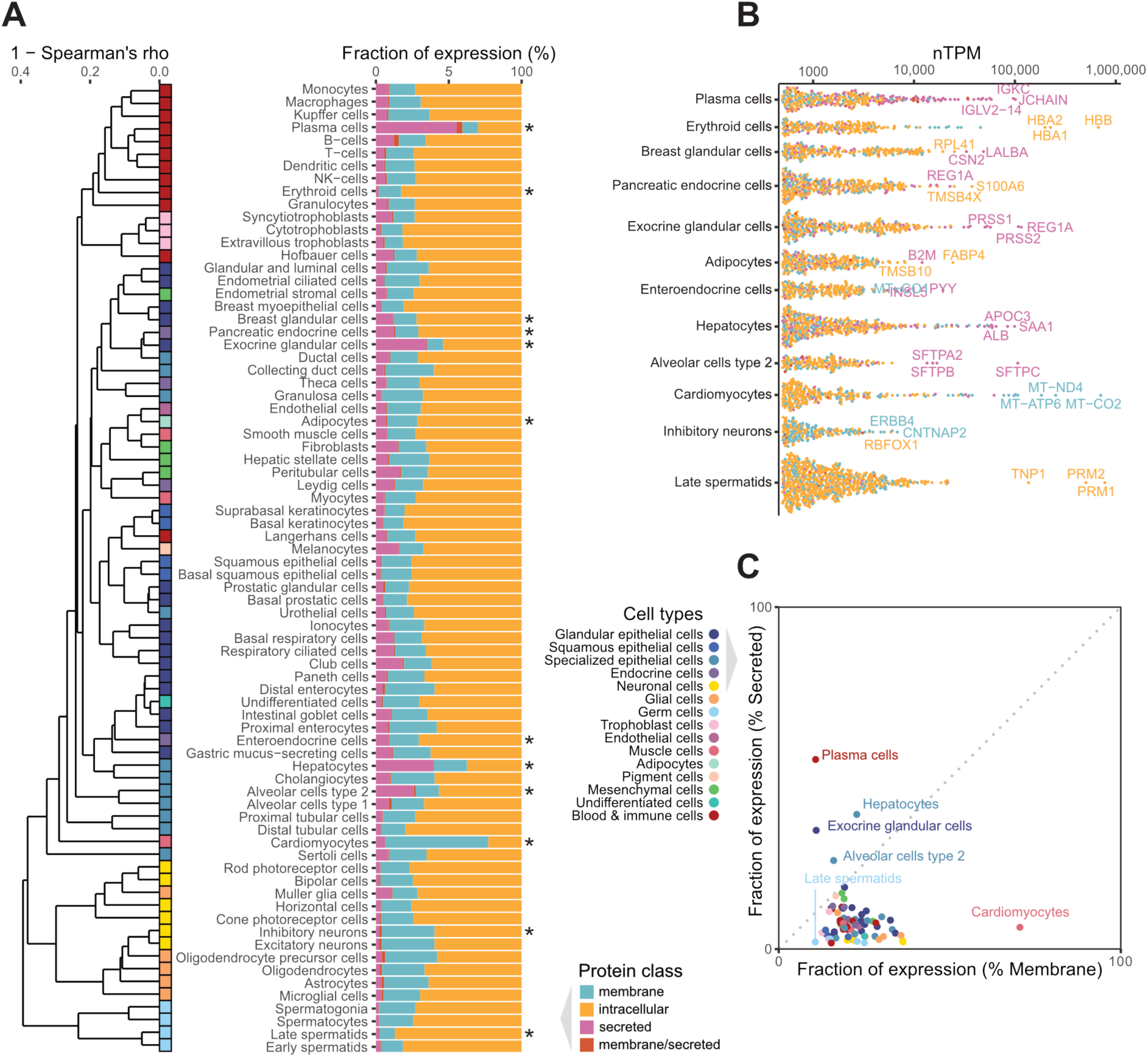
Characterization of the human secretome and membrane proteome across all major cell types. (**A**) Barplot showing the fraction of total gene expression in each single cell type corresponding to genes predicted to encode membrane, intracellular, secreted or membrane/secreted proteins, respectively. The dendrogram is based on the average distance (1 – Spearman’s rho) between the 76 single cell types. (**B**) Beeswarm plot showing the most highly abundant genes (nTPM > 500) in selected cell types based on expression levels, colored by the protein class. Top three genes with the highest expression are labelled for each cell type. (**C**) Relationship between the fraction of total gene expression corresponding to secreted and membrane protein encoding genes in the 76 unique cell types.

On the other side of the spectrum, cardiomyocytes predominantly produce membrane proteins (70%), which is explained by the large amounts of membrane-bound mitochondrial proteins necessary for the high metabolism required for cellular respiration of the heart. The mitochondrial gene Cytochrome c oxidase subunit 2 (MT-CO2) alone constitutes 25% of the total expression of the cardiomyocytes, supporting its important role for generation of energy in the mitochondria. In **Fig. 2B** **and Fig. S4**, the subcellular location of highly expressed proteins is shown, highlighting the top three most highly expressed proteins. The analysis shows that male germ cells have high expression of a large number of genes mainly encoding intracellular proteins. Similarly, the most highly expressed genes in cardiomyocytes are intracellular and relate to mitochondrial function. In contrast, the majority of the highly expressed proteins in plasma cells, pancreatic endocrine cells, hepatocytes and breast glandular cells are secreted. Many of the most highly expressed genes in these cells correspond to well-known secreted proteins, such as IgG related genes in plasma cells, peptidase inhibitor in pancreatic endocrine cells, albumin in hepatocytes and lactoalbumin in breast glandular cells. **Fig. 2C** summarizes the fraction of secreted proteins as compared to the fraction of membrane proteins, highlighting specific cell types such as cardiomyocytes and plasma cells.

The same approach to analyze the fraction of intracellular, secreted or membrane proteins was used for the bulk transcriptomics data. In **Fig. S5A**, the distribution is shown for the 36 tissues. Similar to the results from the single cell transcriptomics analysis, the main outliers include tissues specialized in secretion such as pancreas and salivary gland, whose gene expression consist of a majority of secreted proteins, and heart muscle consisting mainly of membrane proteins (**Fig. S5B**).

### The single cell type specificity of all human genes

A tissue specificity classification based on bulk transcriptomics has been previously described (*13, 14, 36*), allowing all protein-coding genes to be categorized according to their specific expression profiles across all tissues and organs. To complement this analysis and allow for further investigation of cell types present in many tissues, we decided to utilize a similar specificity classification for single cell types using the single cell transcriptomics data described here. Each of the 20,090 human protein-coding genes was classified with regards to their mRNA expression profiles across the 76 single cell types using the criteria outlined earlier and summarized in **Table S1**. 1,908 genes (5%) were categorized as “cell type enriched”, showing high specificity in a single cell type, and 2,799 as “cell type group enriched”, showing high enrichment in a group of up to 10 cell types. In addition, almost half of the genes (n=10,610) were “cell type enhanced”, showing moderately higher expression in a group of 1 to 10 cell types, and 3,260 genes showed “low cell type specificity”, corresponding to 16% of the assessed genes. This group is likely to contain, but is not limited to, many of the “housekeeping” genes necessary for basic cellular functions. Genes with an expression below detection limit (nTPM <1) in all cell types were categorized as “not expressed”, corresponding to 15% of all genes (n = 1,513). Thus, 18,577 genes were expressed in at least one cell type analyzed here. A summary of the number of genes in each specificity category for the single cell transcriptomics analysis is shown in **Fig. 3A**.

**Figure 3.**
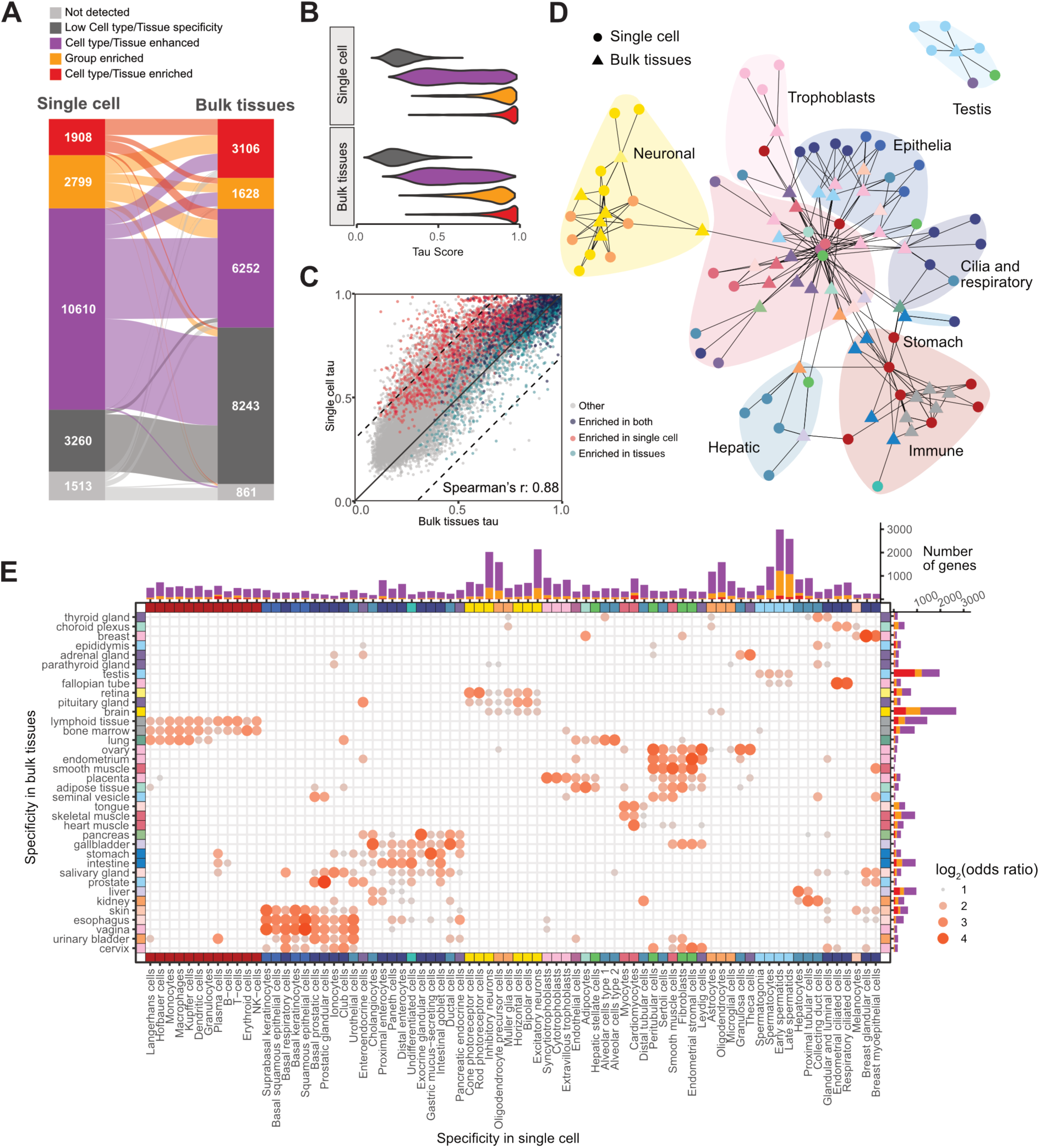
Characterization of gene specificity across single cell types and tissues. (**A**) Alluvial diagram comparing the specificity classification for each category between the single cell (left) and bulk tissue (right) transcriptomics datasets. (**B**) Violin plot showing the distribution of tau specificity score for genes in each of the specificity categories in the single cell dataset (top) and bulk transcriptomics dataset (bottom). (**C**) Scatterplot comparing the tau score in single cell and bulk transcriptomics datasets for each gene. The color code highlights which genes are cell type/tissue or group enriched in both datasets (blue), only in the single cell dataset (red), only in the bulk transcriptomics dataset (cyan), or not enriched in single cell or bulk transcriptomics datasets (grey) (**D**) Network showing top three Spearman correlations between single cells and tissues, calculated based on genes with cell type/tissue or group enriched expression in at least one single cell type or bulk tissue. Regions of the network with related cell types and bulk tissues have been given a general annotation, but an extended version including all cell type and tissue labels can be found in Fig. S9. (**E**) Bubble heatmap showing the overlap between the specificity classification in the single cell and bulk transcriptomics datasets. The bubbles represent a significant overlap (odds ratio > 2, adjusted p-value < 0.05, and number of genes ≥ 3) between genes with elevated expression in the different bulk tissues and single cell types based on odds ratio calculations. The horizontal and vertical barplots visualize the distribution of the categories for each single cell type (top) and bulk transcriptomics (right) datasets, respectively, using the same classification scheme and color code as in (A).

In addition to the specificity classification based on single cell transcriptomics, an updated classification was performed for the bulk transcriptomics data, covering 36 different tissue types. By comparing the classification results between single cell and bulk transcriptomics for all genes (**Fig. 3A**), we found that fewer genes obtained low specificity classification using single cell transcriptomics as compared with bulk transcriptomics. This is expected since many cell types are present across multiple tissues, such as immune cells, fibroblasts and endothelial cells.

Similar to previous reports using bulk transcriptomics and single cell transcriptomics (*13, 14, 41, 42*), testicular germ cells express a large number of genes at elevated levels. Late and early spermatids together have more than 1000 genes that exhibit enriched expression in one of these cell types or both, likely due to their specialized role in the spermatogenesis process unique to male reproduction (**Fig. S6**). Another example of a single cell type with a large number of elevated genes is hepatocytes that has 156 cell type enriched genes, with an overrepresentation of genes related to liver specific functions such as production of proteins involved in complement, activation drug metabolism and hemostasis. Cardiomyocytes have 170 cell type enriched genes, out of which many are involved in muscle structure and contraction. Neuronal and glial cell enriched genes constitute a large number; as exemplified by 90 genes classified as cell type enriched in oligodendrocytes, including myelin oligodendrocyte glycoprotein (MOG) and myelin associated glycoprotein (MAG), and six other genes related to cell adhesion, as well as genes related to ion transport. 310 genes were group enriched in inhibitory and excitatory neurons, including 72 genes related to synapse transmission, assembly, membrane adhesion, and organization. As a collective, neuronal and glial cells constitute the highest number of enriched genes based on specificity classification using the single cell transcriptomics data, with 326 cell type enriched and 1,168 group enriched genes. This supports previous results based on bulk transcriptomics (*13, 36*), suggesting a high number of enriched genes in brain (**Fig. S6**).

### Tau scoring of gene expression for spatial specificity

In addition to annotating each gene with a discrete category denoting the gene’s specificity, a continuous specificity metric score was calculated as tau (*43*), a metric ranging from 0 to 1 in order of increased specificity, where a gene with 0 tau would indicate identical expression across all cells or tissues, and a tau of 1 indicates that the gene is expressed only in a single cell or tissue (*43*). We observed a range of tau from 0.1 to 1.0 across the single cell transcriptomics dataset with high concordance with the specificity categorization scheme (polyserial correlation: 0.83, **Fig. 3B**) with cell type enriched genes having a median tau of 0.90, while the corresponding values for group enriched is 0.86, cell type enhanced 0.63, and low cell type specific genes 0.30. This shows that the two specificity metrics for determining single cell type specificity yield complementary results, which is reassuring since one uses a continuous metrics (tau) and the other uses an arbitrary cut-off (nTPM <1).

We observed that, in general, the same genes had similar specificity classification when comparing the single cell and bulk transcriptomics datasets. The Spearman correlation of tau between the two datasets across all genes was 0.88 (**Fig. 3C**). This demonstrates that the majority of genes with high specificity using single cell transcriptomics show the same specificity classification using bulk transcriptomics (**Fig. S7**). However, 1073 genes showed higher specificity using single cell transcriptomics (**Fig. S8A**). These genes are of particular interest, as they represent expression patterns that are not reflected in bulk tissue data and showcase the additional information brought by single cell-based analysis. An overrepresentation analysis based on cell markers from PanglaoDB (*44*) as well as genes with enriched cell type expression according to our own analysis showed that these genes show high expression in single cell types present in multiple tissues, such as endothelial cells, fibroblasts, pericytes, smooth muscle cells, adipose cells, stromal cells, and plasma cells, or in single cell types with relatively low abundance in their tissues, whose expression is therefore masked using bulk transcriptomics analysis, including peritubular and Sertoli cells in testis, and glial cells in brain (**Fig. S8B**). An additional analysis of these 1073 genes to Gene Ontology (*45, 46*) indicated similar results, with enrichment of genes related to the extracellular matrix, cell adhesion, angiogenesis, and immune response (**Fig. S8C**).

### Comparison between single cell transcriptomics and bulk transcriptomics

While a high correlation between single cell and bulk transcriptomics analysis based on tau demonstrates that gene specificity is highly similar between the two datasets for the majority of genes, the analysis does not take into consideration which tissues and single cell types the genes show enrichment within. To visualize the correlation between single cell and bulk transcriptomics data for genes with elevated expression, a network was constructed connecting each cell type in the single cell transcriptomics dataset with the highest correlating tissue in the bulk transcriptomics dataset (**Fig. 3D** **and Fig. S9**). The results for all genes are visualized in a heatmap (**Fig. S10**) demonstrating that the bulk tissue data in general shows close correlation with single cell type data across all the human major tissues and organs.

To further investigate similarities in gene specificity between single cell and bulk transcriptomics and to disentangle which cells and tissues that share an overlap of co-enriched genes, a one-sided hypergeometric test was performed to assess the pairwise overlap between cell type enriched genes and tissue enriched genes. The result is displayed in a bubble heatmap (**Fig. 3E**), where each bubble corresponds to a statistically significant overlap. This plot also displays the number of genes in each of the specificity classification categories per single cell type and tissue, respectively. It is reassuring that single cell types specific to a certain tissue such as cardiomyocytes (heart), germ cells (testis), and trophoblasts (placenta) analyzed with single cell transcriptomics show high overlap with the corresponding classification based on bulk transcriptomics. Cells with similar function also overlap, such as urothelial cell enriched genes with those that have enriched expression in urinary bladder or squamous epithelia, including esophagus. Hepatocytes overlap primarily with liver tissue, but some overlap can also be observed with kidney tissue which shares many group-enriched genes with liver. Cardiomyocytes overlap with bulk transcriptomics analysis of heart muscle, skeletal muscle and tongue, all containing striated muscle cells, while myocytes overlap with skeletal muscle and tongue tissue, showing that despite many similarities between different striated muscle cells, myocytes and cardiomyocytes still have unique expression profiles.

Several epithelial cell types (basal squamous epithelial cells, suprabasal keratinocytes, basal keratinocytes, and squamous epithelial cells) show an overlap in expression to their corresponding tissues (vagina, esophagus, and skin), with additional overlap between single cell types and tissues that have similar types of cells or functions. Such similarities that do not directly correspond to the tissue of origin for specific single cell types include for example basal prostatic cells and seminal vesicle, as well as several different glandular cell types that showed an overlap both to their tissue of origin and other tissues containing glandular cells. Another example is enteroendocrine cells that showed an overlap to both gastrointestinal tissues known to include enteroendocrine cells, but also other organs with endocrine functions such as pancreas, pituitary, and adrenal gland.

### New classification of all protein-coding genes based on expression clusters

The specificity classification based on single cell and bulk transcriptomics (*13, 36*) is helpful for stratifying genes based on elevated expression in any cell type or tissue. However, many important groups of genes share more subtle expression patterns that will be missed in this specificity classification scheme. One example includes ribosomal genes, which are co-expressed and form a distinct expression pattern between single cell types, but are expressed ubiquitously with low specificity, which means that these genes will be lumped together with all other genes with low specificity. To extend the annotation of expression patterns across genes, gene clustering can be devised to stratify genes into distinct clusters of similarly expressed genes, whether the expression patterns show high specificity or not, potentially highlighting previously unknown relationships between genes.

A novel gene clustering analysis was therefore performed using both the single cell transcriptomics data and bulk tissue transcriptomics data utilizing the pipeline as outlined in **Fig. 4A**, including gene clustering and subsequent manual cluster annotation. The final consensus clustering results consisted of 89 tissue gene expression clusters based on the bulk transcriptomics dataset (**Fig. 4B**, **Fig. S11**) and 68 single cell type gene expression clusters based on the single cell transcriptomics dataset (**Fig. 4C**). In addition, two separate cluster analyses were performed for expression across 18 flow sorted immune cell populations and across 69 cell lines (data not shown, but available at www.proteinatlas.org).

**Figure 4.**
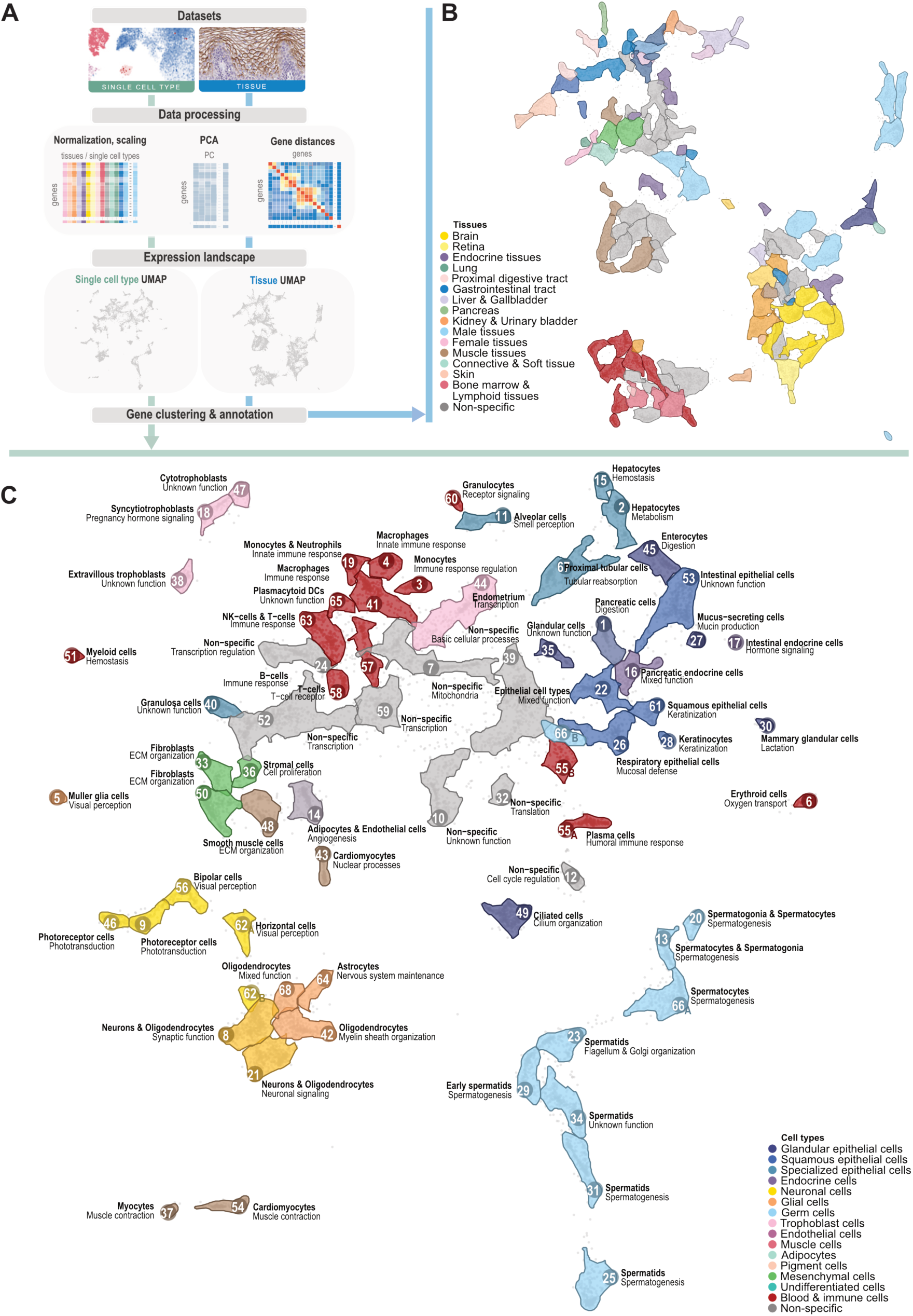
Clustering of protein-coding genes based on expression across single cell types and bulk tissues. **(A)** Overview of the pipeline used to cluster the protein-coding genes, starting with the bulk transcriptomics and single cell HPA datasets, respectively, followed by pre-processing of the data, visualization of the gene expression landscape in a UMAP, and gene clustering based on the expression profiles across bulk tissues and single cell types. The resulting clusters are manually annotated in terms of specificity and function and visualized in the UMAP accordingly. **(B)** UMAP visualization of the bulk transcriptomics tissue expression clusters. The cluster colors are assigned based on the annotated specificity. The individual colors are mixed in case of multiple tissues. An extended version including the specificity and functional annotation is included in Supplementary Fig. S14. **(C)** UMAP visualization of the single cell expression clusters. Clusters are colored according to the single cell type specificity of the cluster and labelled according to the annotated specificity (bold) and function. labelled according to the annotated specificity (bold) and function.

In the UMAP visualization (**Fig. 4C**), each cluster was colored based on cell or tissue type specificity of the cluster genes. Although cluster annotation was performed independently from the UMAP visualization, clusters with similar specificity and related function reside close together in the UMAP visualization. Interestingly, the non-specific clusters reside at the center of the UMAP (**Fig. 4C** and **Fig. S12-14**) and have annotated functions related to typical housekeeping functions such as “Basic cellular processes”, “Mitochondria”, “Transcription”, “Transcription regulation”, and “Translation”. Immune related clusters mostly group closely together, with monocytes, macrophages, plasmacytoid dendritic cells, granulocytes, T-cells, B-cells, and plasma cells, each represented by their own cluster. Cluster 54 “Cardiomyocytes - Muscle contraction” with 250 genes reside as expected in close proximity to Cluster 37 “Myocytes – Muscle contraction”, while the other heart muscle cluster 43 “Cardiomyocytes - Nuclear processes” with 192 genes is closer to other clusters with housekeeping functions.

### Genome-wide annotation of all protein-coding genes based on expression clusters

Each gene cluster was manually annotated by experts using a multitude of external and internal datasets and predictions in terms of gene function and protein localization, as well as cell and tissue type specificity of the genes present in each cluster. A summary is shown in **Fig. 5** for the single cell transcriptomics dataset and **Fig. S15** for the bulk transcriptomics dataset. The characteristics of all genes in a cluster are shown including the total number of genes, the specificity score (tau), the cell type specificity (enriched, enhanced or low specificity). In the final consensus clustering of 68 single cell type gene expression clusters based on single cell transcriptomics, 61 were annotated to show specificity for a number of single cell types while seven were annotated as “non-specific”, and 57 were annotated as having a main function (**Fig. 5** and **Supplementary data S4**). In **Fig. 5** (bottom), the average expression of genes in each cluster across single cell types is shown. The heatmap demonstrates that most of the clusters contain genes with elevated expression in a particular cell type, as exemplified by hepatocyte-related genes in cluster 2 “Hepatocytes – Metabolism” and cardiomyocyte-related genes in cluster 54 “Cardiomyocytes - Muscle contraction”. In contrast, several gene clusters show ubiquitous expression across the cell types and these clusters are therefore annotated as “non-specific” harboring genes with essential functions, such as transcription and translation.

**Figure 5.**
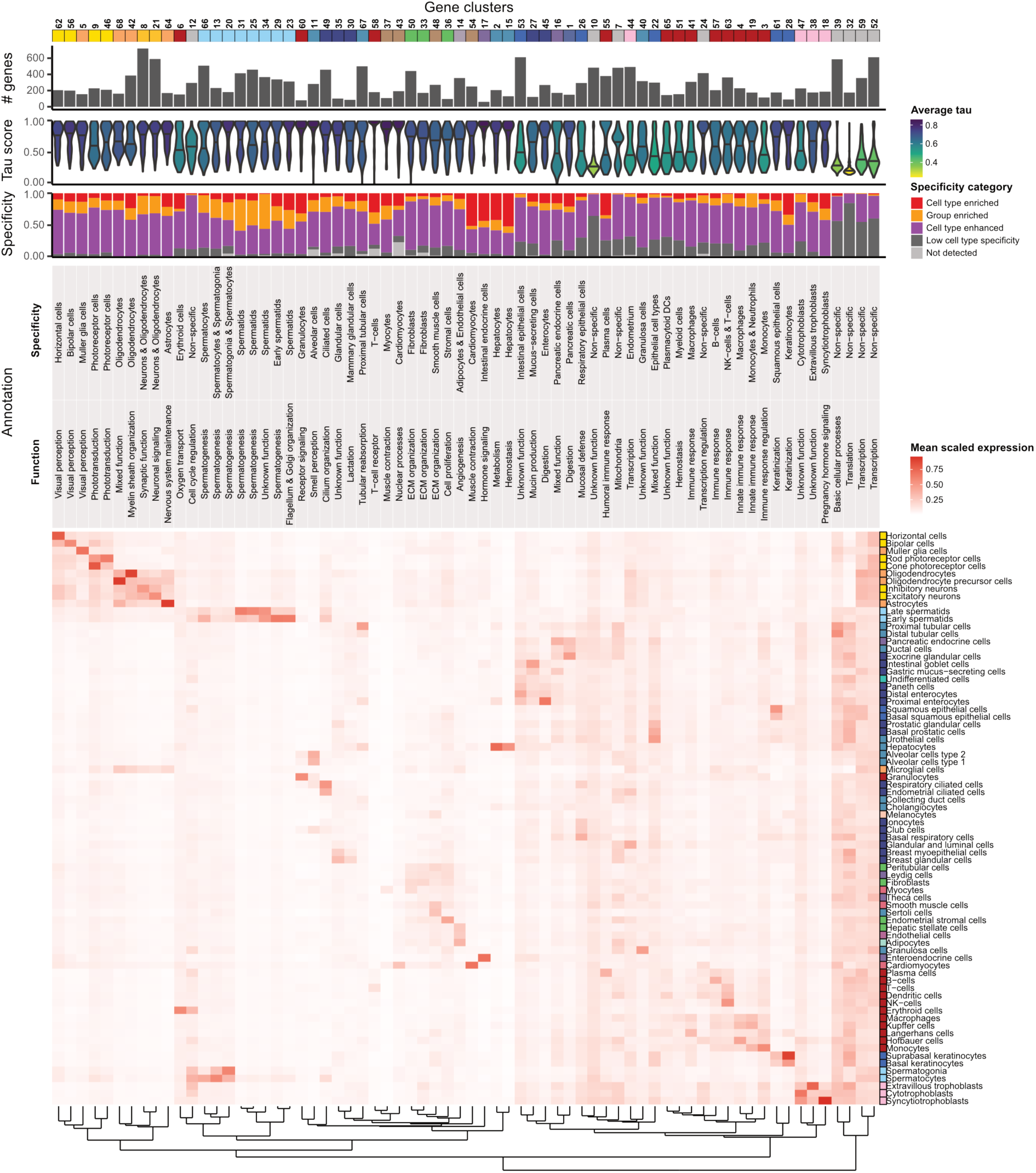
Characterization of single cell gene expression clusters. The number of genes, distribution of tau scores, proportion of genes in each specificity category, annotated specificity and function and summarized mean gene expression across the 76 cell types is indicated for each of the 68 clusters. The dendrogram is obtained by hierarchical clustering of distances based on gene expression levels for all single cell types using Ward’s criterion.

### Annotation of the ciliated cell types

The analysis of gene clusters allows for exploration of uncharacterized proteins that are present within a certain cluster. In several cases, subtle, yet distinct, patterns were captured, where genes related to a specific function formed a cluster. This is the case for single cell type cluster 49 – “Ciliated cells - Cilium organization”, which contains 457 genes mainly highly expressed in ciliated cells or cells containing flagella (respiratory ciliated cells, endometrial ciliated cells, spermatocytes, early- and late spermatids). A more in-depth analysis showed that the genes in this cluster had a large overrepresentation of genes annotated as part of cilium, as well as many related terms (“cilium assembly”, “cell projection”, “axoneme”, etc). In addition, the cluster showed significant overlap to PanglaoDB cell type markers for ependymal cells, olfactory epithelial cells, ciliated cells, and germ cells, as well as an overrepresentation for genes with elevated expression in fallopian tube, choroid plexus, and testis based on bulk transcriptomics analysis, representing tissues rich in ciliated/flagellated cells.

The corresponding proteins in cluster 49 were further validated by antibody-based profiling using human tissue samples, including airway epithelia (nasopharynx and bronchus), fallopian tube and testis. Based on a stringent validation pipeline, reliable protein expression could be confirmed in ciliated cells, flagella or a combination thereof for 300 out of the 457 genes in the cluster (**Fig. 6A** **and Supplementary data S5**) supporting the cilia-related function for these genes in cluster 49 by bioimaging. Interestingly, the cell type specificity for these 300 proteins formed two main groups: proteins expressed only in ciliated cells of airway epithelia and fallopian tube (n = 179), and proteins expressed both in ciliated cells of airway epithelia and fallopian tube as well as in sperm flagella (n = 84) (**Fig. S16**). This suggests that despite cilia and flagella sharing a common basic structure and being the fundamental units of motion in cellular biology, ciliated cells maintain a distinct protein expression profile. This is not unexpected since cilia are generally shorter than flagella and they are present on the cell surface in much greater numbers, with up to hundreds of cilia on a single cell, while flagellated cells usually have a single flagellum. The in-depth antibody-based profiling analysis suggests spatial expression forming four main groups of expression related to cilia/flagella, including (i) proteins expressed only intracellularly in ciliated cells (n = 54), (ii) proteins expressed only in the cilia axoneme (n = 47), and (iii) proteins expressed in both intracellularly and in the cilia axoneme (n = 104), as well as proteins expressed intracellularly and in the cilia axoneme of ciliated cells combined with expression in sperm flagella (n = 57) (**Fig. 6A**). One example of a protein distinctly expressed both in the axoneme tip of ciliated cells in sperm flagella, suggesting a function related to motility in humans is the uncharacterized protein C4orf47 (**Fig. 6A**). Other examples of previously characterized proteins include C1orf87, observed on the tip of cilia and ciliary rootlets in fallopian tube and airway epithelia (**Fig. S17**) and C20orf85 expressed in cilia axoneme, tip and rootlets, previously found to be under-expressed in respiratory cilia of children with severe asthma (*47*) (**Fig. S17**).

**Figure 6.**
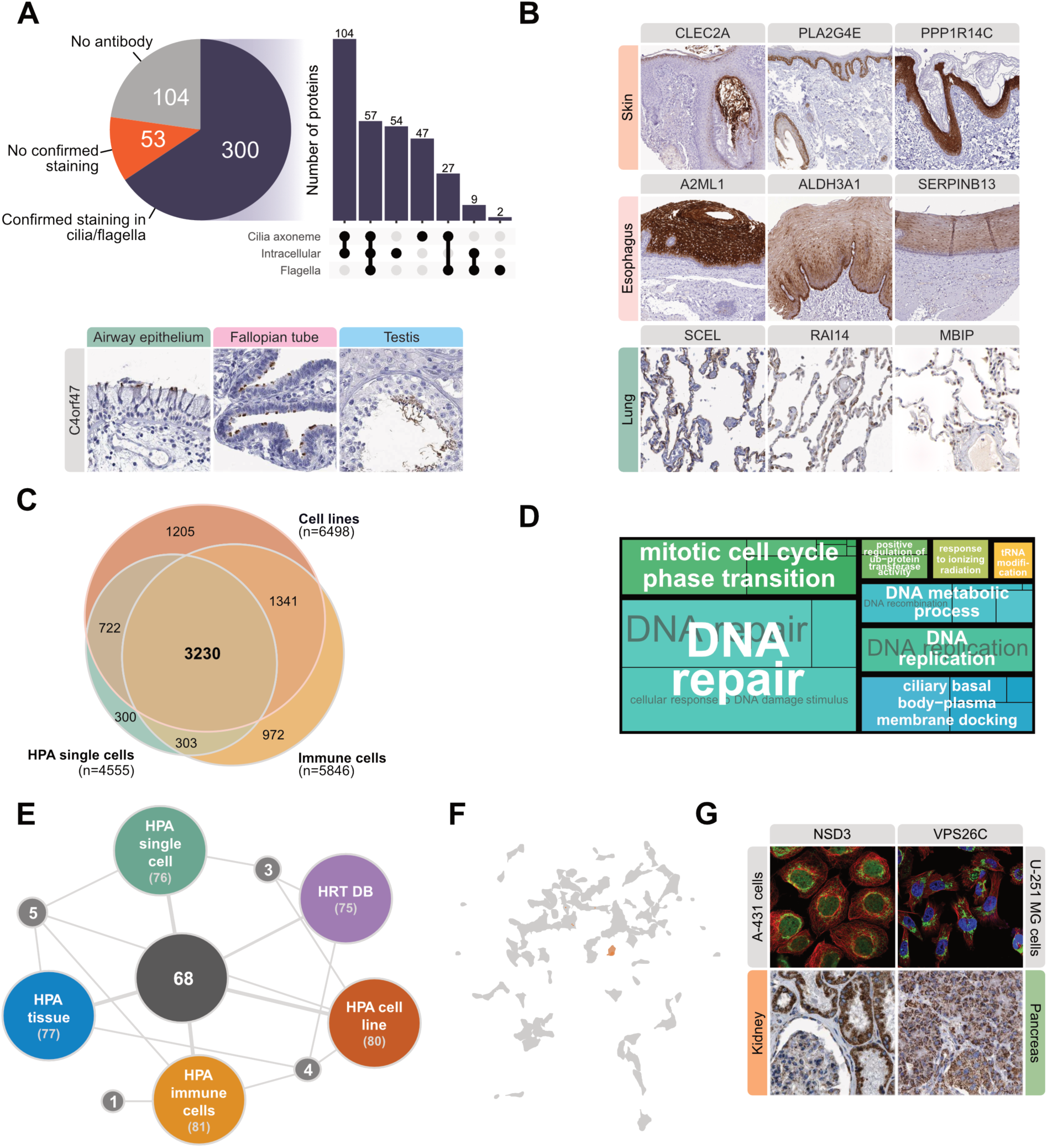
Exploration of cell type specific and housekeeping protein classes. (**A**) Summary of IHC staining pattern for proteins in cluster 49 in the single cell type gene-based clustering results. Proteins are classified into three categories: (i) confirmed staining in cilia/flagella, (ii) no confirmed staining, and (iii) no antibody (top). IHC images in airway epithelium, fallopian tube and testis for the C4orf47 protein. The upset plot shows the staining pattern in cilia axoneme, intracellular space and flagella for proteins with confirmed staining in cilia/flagella. (**B**) Examples of previously unknown or less-well studied proteins with respect to biology of skin (top row), esophagus (middle row) and alveolar epithelial cells (bottom row) stained with immunohistochemistry. (**C**) Overlap of genes identified as housekeeping in the single cell type, cell line and immune cell HPA datasets, respectively. (**D**) Treemap summarizing the overrepresented GO terms (adjusted p-value < 0.01) related to the Biological Process Ontology for the 1205 genes expressed only in the cell line dataset. (**E**) Summary of the overlap between genes annotated as ribosomal in different datasets. HRT DB = Housekeeping Transcript Atlas database. (**F**) Single cell type gene expression UMAP highlighting the single cell gene expression cluster with an overrepresentation of ribosomal genes. All ribosomal genes are colored in orange. (**G**) IF and IHC images show the location of example proteins NSD3 (in A-431 cell line and kidney) and VPS26C (in U-251 MG cells and pancreas).

Another example from this cluster is Androglobin (ADGB), a protein named after its high expression in testis (*48*), but lacks annotations related to cilia in all functional databases assessed (GO, KEGG, and Reactome). Our analysis shows an expression profile consistent with that of a cilium related gene, confirmed by antibody-based profiling displaying protein localization restricted to the tip of cilia the cilia axoneme in both airway epithelia and fallopian tube (**Fig. S17**). The presence of ADGB in ciliated cells was further supported by co-expression with two transcriptional regulators shown to regulate the expression of ADGB: Forkhead box J1 (FOXJ1) (*49*), an important transcription factor in ciliogenesis (*50*), and Regulatory factor X2 (RFX2), a major transcriptional regulator in spermiogenesis (*51*). Interestingly, both FOXJ1 and RFX2 are also members of the expression cluster 49 together with ADGB. While the exact role of ADGB in cilia is yet to be determined, it has been speculated to contribute to cilia biogenesis and function (*49*). In summary, using the new annotation tool based on genome-wide annotation of the protein-coding genes based on spatial expression clustering, it was possible to perform a comprehensive annotation of proteins expressed in ciliated cells and proteins simultaneously expressed in flagella, as well as proteins expressed in different structures of the ciliated cells and thus contributing to the functional annotation of proteins expressed in ciliated cell types.

### Annotation of the skin-related proteome

Another example of an interesting gene cluster is cluster 28 “Keratinocytes – Keratinization”, which includes 89 genes with a large proportion of well-known proteins associated with epidermal keratinocytes and keratinization, but also proteins with unknown expression patterns and/or functions. An example of such protein is CLEC2A, a C-type lectin-like receptor implied in the recognition, entrapment, and spreading of HIV and Dengue virus upon platelet activation, as well as in the enhancement of immune response and organ damage in viral infections (*52*). The presence of this protein in cluster 28 suggests a skin-related function and antibody-based profiling confirms expression of CLEC2A in granular and corneal layers of normal epidermis (**Fig. 6B**), suggesting a role in skin barrier function. PLA2G4E is a phospholipase with a proinflammatory function and its expression has been shown to be absent in normal skin, but increased in the upper epidermal layers in patients with psoriasis and pityriasis rubra pillaris (*53*). Antibody-based profiling of PLA2G4E (**Fig. 6B**) shows localization of the protein in basal keratinocytes, supporting a skin-related function and calls for in-depth analysis to explore its role in normal and diseased skin. Another protein in this cluster is PPP1R14C, a protein phosphatase inhibitor, which is poorly characterized in human tissues. It is present in brain, heart, and skeletal muscle, where its expression is regulated by acute and chronic morphine treatments (*54*) but its role in skin is unknown. The antibody-based profiling (**Fig. 6B**) shows cytoplasmic expression throughout all layers of the epidermis, thus supporting a skin-related function.

Another skin-related cluster is 61 “Squamous epithelial cells – Keratinization” which contains 176 genes, including several poorly characterized proteins. The protein A2ML1 was initially shown to be a protease inhibitor expressed in keratinosomes of granular layer in the skin and secreted extracellularly in stratum corneum, with a potential role in skin barrier function and desquamation (*55*). It was also identified as p170 antigen recognized by autoantibodies in paraneoplastic pemphigus (*56*), an autoimmune blistering disease that affects squamous epithelia (*57*). Antibody-based profiling shows expression of A2ML1 in suprabasal layers of esophagus (**Fig. 6B**), supporting its role in the development of esophageal lesions. The function of A2ML1 in tissues with mucosal squamous epithelia is unknown, although gene expression has been shown to be downregulated in squamous cell carcinoma of the esophagus (*58*). In addition, the protein ALDH3A1 is one of the main proteins in cornea which plays a protective role against oxidative stress, including UV-induced oxidative stress (*59*). Previous reports of its expression in normal and neoplastically transformed epithelia were conflicting (*60, 61*).

Antibody-based profiling showed a consistent expression profile with the cluster analysis, localized to the epithelium of esophagus (**Fig. 6B**), suggesting a possible protective role against oxidative stress in the esophagus. In addition, the protein SERPINB13 is a skin serin-protease inhibitor, which inhibits Cathepsine L and protects keratinocytes against UV-induced apoptosis. In normal epidermis, it is expressed in the basal layer, while the protein is redistributed to upper layers upon inflammatory disease. Here, the antibody-based profiling showed distinct expression of SERPINB13 in suprabasal squamous epithelial cells of the esophagus (**Fig. 6B**), suggesting that this protein also has other functions. These examples again demonstrate the value of the new genome-wide annotation strategy to identify proteins for more in-depth functional studies.

### Annotation of the alveolar proteome

Another interesting expression cluster is the 281 genes in cluster 11 “Alveolar cells – Smell perception”. Several of these proteins are known to be associated with alveolar cells and/or smell perception, but also contains several proteins with unknown expression patterns and functions in alveoli and no known association to gas exchange or surfactant homeostasis. The protein SCEL was discovered as a component of cornified envelope in keratinocytes, squamous epithelia and in elastic and muscle arteries suggesting a role in resilience to biomechanical stress in these tissues (*62, 63*). Using antibody-based profiling, we here show highly enriched expression of SCEL in type 1 alveolar epithelial cells (**Fig. 6B**) suggesting a role of mechanical plasticity of alveolar cells. The protein RAI14 is a polyglutamine encoding gene expressed in several tissues, especially during the development (*64*). It is functionally associated with cortical actin-cytoskeleton and testis-specific actin-rich adherens junctions and plays a role in membrane shaping during neuron development (*65, 66*). The functional relevance of RAI14 for the biology of alveolar epithelial cells is so far unknown, but here we demonstrate with antibody-based profiling that this protein is specifically expressed in alveolar epithelial cells type 2 (**Fig. 6B**), indicating a role in surfactant secretion. Another interesting protein with nuclear expression in both types of alveolar cells was MBIP, a leucine zipper-like motif-containing protein encoded by a gene located in close proximity to NKX2-1. Deletion of both genes is detrimental and causes Brain-lung-thyroid syndrome characterized by congenital hypothyroidism and infant respiratory distress syndrome, suggesting their role in the development of lungs and thyroid gland (*67*). Amplification of the MBIP gene is a frequent event in lung carcinoma and its expression is associated with worse survival of patients with lung carcinoma (*68*), however the expression pattern and function of MBIP in adult lung is unclear. Here, we show with antibody-based profiling (**Fig. 6B**) that this protein shows distinct location to the alveolar cells of the lung.

### Integration of gene clustering data

An overrepresentation analysis using a one-sided hypergeometric test was performed to make a pairwise comparison between clusters based on the single cell transcriptomics and bulk transcriptomics analyses to assess whether the clusters shared more genes than expected by chance. Interestingly, many independent clusters shared an overrepresentation of genes, and these clusters often had annotations of similar specificity and functionality. The result is displayed in a network (**Fig. S18**) where each node signifies a cluster, and each connection represents a statistically significant overlap of genes. As an example, the bulk tissue transcriptomics expression cluster 13 and 33 “Striated muscle – muscle contraction” overlaps with the single cell transcriptomics cluster 37 “Myocytes – muscle contraction”, and 54 “Cardiomyocytes – Muscle contraction”, which in turn is overlapping with two bulk tissue gene clusters with annotations related to heart muscle contraction. Similar connections between tissues and the corresponding single cell types unique for a certain tissue type are observed for immune-related tissues, pancreas, kidney, liver, intestine, skin, retina, brain and testis. Tissues and single cell types that are closely related also cluster in proximity, such as retina and brain. Interestingly, tissues and single cell types with a secretory function, such as mucus-secreting cells, mammary gland, breast and salivary gland have a clear connection. Pituitary gland, that carries out both neuronal and endocrine functions, clusters between brain related tissues and single cell types and intestinal endocrine cells, thereby representing both functions. In conclusion, it is reassuring that the new genome-wide annotation yields complementary validation using the two complementary datasets, single cell and bulk transcriptomics, respectively.

### The essential (“house-keeping”) proteome

Housekeeping (HK) genes are constitutively expressed genes that are required for the maintenance of basic cellular functions. Despite their importance in the calibration of gene expression, as well as the understanding of many genomic and evolutionary features, a single definition does not exist, and parameters such as expression variation and gene or transcript level data are sometimes, but not always, included, and important discrepancies have been observed in studies that previously identified these genes. The bulk transcriptomics analysis in the HPA Tissue Atlas from 2015 (*13*) showed that close to 9000 genes were expressed in all tissues, suggesting that these genes are needed in all cells to maintain basic cellular structure and function. This is a considerably higher number than the suggestion from Eisenberg (*69*), suggesting that 3,804 genes encode housekeeping proteins. Recently (*70*), these studies have been updated by the Housekeeping Transcript Atlas database (HRT) using bulk transcriptomics data from the GTEx consortium (*5*) and thus redefining human housekeeping genes and candidate reference transcripts by mining massive RNA-seq datasets. This study identified 2176 genes expressed in all 52 tissues analyzed. Here, we decided to investigate the set of housekeeping genes based on the single cell transcriptomics dataset resulting in the identification of 4555 genes expressed above detection cutoff (≥1 nTPM) in all analyzed single cell types (**Fig. 6C**). These numbers are considerably larger than the HRT study (n = 2176), which is likely due to the fact that we have not applied any criteria for expression variance. To extend this analysis, we included data from human cell lines and immune cells sorted by FACS (*36*) and compared the results from the single cell transcriptomics dataset. The analysis (**Fig. 6C**) showed that 3,230 genes were identified as expressed in all samples in all three separate datasets. A functional analysis revealed (**Fig. S19A**) that among these 3230 genes are many genes belonging to known housekeeping functions, such as translation and gene expression. Many genes were found to be expressed in all cell lines, but not in the immune cells and the single cell types present in tissues. A functional analysis of these 1205 genes expressed in all cell lines (**Fig. 6D**) shows that many of these genes are related to cell cycle control and DNA repair supporting a functional role for rapidly dividing cells. This is exemplified by the protein Golgin A5, for which immunohistochemical analysis showed Golgi-like expression in all analyzed single cell types (**Fig. S19B**). In this context it is interesting that only 20% of the proposed housekeeping genes have so far been established as essential using CRISPR knock-out data in the DepMap portal (*71, 72*) and the extended list of proteins suggested to be housekeeping in our study must therefore be validated by other approaches to establish their role as essential in human cells.

An important group of essential genes is mitochondrial genes, constituted by 13 mitochondrial encoded and 1231 nuclear encoded genes according to MitoCarta3.0 (*73*), while in UniProt the number of reviewed genes with subcellular location mitochondrion is 1270 (*74*), whereas 947 genes are overlapping the datasets. The cluster analysis reveals that all 13 of the mitochondrial encoded genes reside in cluster 54 “Cardiomyocytes – Muscle contraction”, while nuclear encoded genes have a disperse spatial distribution (**Fig. S20**), with a statistically significant overrepresentation of mitochondrial genes in clusters 7, 39, 54 and 67. Cluster 7 “Non-specific – Mitochondria” 7 contains 481 genes, out of which 60 are listed as mitochondrial genes by MitoCarta. Interestingly, several of the genes not annotated as mitochondrial in this cluster show a mitochondrial pattern using the antibody-based subcellular profiling as demonstrated by the bioimaging from the HPA shown in **Fig. 6G**. The antibody-based profiling confirm the mitochondrial staining both in tissues and cell lines, exemplified by the methyl transferase Nuclear receptor binding SET domain protein 3 (NSD3), and VPS26 endosomal protein sorting factor C (VPS26C). While VPS26C has a suggested function in endosomes, neither NSD3 nor VPS26C have previously been described in the context of mitochondria. These results demonstrate the usefulness of the genome-wide annotation using expression clusters to identify mitochondria-related genes/proteins that should be further explored.

Another interesting group of housekeeping genes are those encoding ribosomal proteins. These include proteins that, in conjunction with rRNA constitute ribosomal subunits involved in translation. Altogether, 82 protein-coding genes assemble the human ribosomes, and 80 of these are highly essential according to CRISPR knock-out data in the DepMap portal (*71*). **Fig. 6E**, shows the expression of these ribosomal genes in different datasets, showing that the great majority of ribosomal genes are ubiquitously expressed and classified as “housekeeping” in the HRT (*70*). Most of the ribosomal genes (n = 78) are found in one gene cluster (32 “Non-specific – Translation”) suggesting a similar expression in all single cell types (**Fig. 6F**), while four are found in other clusters. One of these is ribosomal protein S4 Y-linked 2 (RPS4Y2) located on the Y chromosome, and has, as expected, been suggested as non-essential in humans (*75*) and this protein has also been found non-essential in the DepMap portal, supporting the annotation of this ribosomal protein as not belonging to the housekeeping gene set. The cluster 32 “Non-specific – Translation” constitutes 175 genes in total, and thus also contains many non-ribosomal genes that share a similar expression profile to the ribosomal protein encoding genes. A noteworthy example is the Tumor protein, translationally-controlled 1 (TPT1), which has independently been found to interact with ribosomal proteins in the Open Cell program (*10*). This demonstrates that the cluster analysis, by capturing similarities in gene expression profile also can be used to find potential interaction partners, which can be further validated by more in-depth studies.

### The druggable proteome

Almost all pharmaceutical drugs act by targeting proteins and modulating their activity (*13*). According to DrugBank (*76*), >1,000 protein targets are approved for drugs in the US targeting enzymes, transporters, ion channels and receptors, respectively. Of these targets, 807 are represented in our dataset, and the UMAP cluster analysis (**Fig. S21A**) shows that the genes encoding drug targets are scattered across the gene clusters, which is consistent with pharmacological drugs targeting proteins with different expression profiles across single cell types. An important protein family is the G-protein coupled receptors (GPCR) that make up more than 10% of the pharmacological targets for FDA approved drugs (*77*). The family can be divided into the 371 olfactory receptors involved in smell and taste and the 393 non-olfactory receptors which include many drug targets, having multiple important biological roles in the human body. Changes in receptor activity have been shown in many disorders, including tumorigenesis, and drugs targeting GPCRs are used for treatment of pain, inflammation, neurobiological, metabolic disorders (*78*). The UMAP analysis (**Fig. S21B**) reveals that the non-olfactory GPCRs are widely, but not evenly distributed across the clusters with the most significant overrepresentation found in brain and neuronal-related clusters 8 and 21, visual perception cluster 62, ECM organization cluster 48 and immune related clusters 63 and 19. The olfactory GPCRs are less widely distributed, but they are also less represented, since more than 60% are not expressed in the analysed single cell types and therefore not part of the analysis. The clusters with most significant overrepresentation of olfactory receptors include smell perception cluster 11, neuronal signalling cluster 21 and granulocyte cluster 60 (**Fig. S21C**). An example of a drug target belonging to the GPCR family is Calcium sensing receptor (CASR), which plays a key role in maintaining calcium homeostasis. By activating calcium-sensing receptors on parathyroid cells, secretion of the parathyroid hormone is suppressed, playing an important role in the treatment of hyperparathyroidism. The antibody-based profiling supports a high membranous expression in the parathyroid gland (**Fig. S21D**).

### Tissue-specific cell types

In this study, the analysis has been performed on the 76 consensus cell types in which identically annotated cell types from different tissues have been combined. However, the data also allows us to probe the different expression profiles of the same cell type across different tissues. Some of the cell types are indeed found in a large number of tissues and organs, including T-cells in 17 tissues, macrophages in 15 tissues, fibroblasts and smooth muscle cells in 11 tissues and endothelial cells in 14 tissues (**Fig. S1B**). An analysis of the difference among 24 clusters annotated as endothelial cell types reveals that those from pancreas, lung and heart are outliers in a global analysis using PCA dimensionality reduction (**Fig. S22A**). Moreover, UMAP dimensionality reduction shows that genes have distinct expression profiles across endothelial clusters, for example higher expression in muscle or lung endothelia (**Fig. S22B**).

This calls for a more in-depth analysis to explore the tissue-specific difference of the same cell type across the analyzed tissues and organs. As an example, a comparative analysis of gene expression in endothelial cell clusters from different tissues revealed that *SELE*, which encodes an adhesion receptor critical for leukocyte recruitment during inflammation (*79*), is expressed at highest levels in skin endothelial cells, compared to those in other tissues, an observation validated by antibody-based profiling (**Fig. S22C**). This observation is consistent with its previously reported role in the constitutive immune surveillance in skin (*80*). Another example, *FCGR2B,* a low affinity receptor for the FC region of immunoglobulin gamma complexes, was markedly elevated in placental endothelial cells, compared to those from other tissues (**Fig. S22C**). This receptor is a well-known marker of liver sinusoidal endothelial cells, where it has a role in the phagocytosis of immune complexes (*81*), but its role in the placental vasculature is less well studied. These examples suggest specialized functional roles of these cell types within different tissues, including a role of regulatory influence of their respective microenvironments.

### The single cell type section of the Human Protein Atlas

A new open access version 21.1 of the single cell section of the Human Protein Atlas has been launched (www.proteinatlas.org). The new section contains results from the 444 single cell type clusters, representing 76 major single cell types across 25 tissues. The resource contains genome-wide data on all 20,090 individual protein-coding genes including the new classification for single cell type specificity called tau. In addition, each protein-coding gene is classified according to spatial expression profiles into altogether 68 manually annotated single cell type expression clusters to define genes with similar expression across major cell types in the human body. Complementary gene expression clustering analyses based on expression based on bulk tissue transcriptomics as well as datasets from cell lines and blood cells are also available on the online portal. For each protein-coding gene, the nearest neighboring genes with regards to spatial expression across the human single cell types are presented, giving information on co-expressed genes to facilitate functional hypothesis generation for genes with incomplete functional annotation.

## Discussion

An important quest for genome annotation research is to define a complete “parts-list” of the building-blocks of human life, the genes and the proteins, and their relationship to human biology and disease. It is in this context important to make the annotation available and interpretable to a wide audience. Here, we have presented a revised annotation of all human protein-coding genes based on RNA and protein profiling across tissues and organs in the human body and with a focus on single cell transcriptomics data. Altogether 25 different tissues and organs have analyzed using a pooling strategy to achieve high quality data also for genes with low expression levels.

The single cell and spatial transcriptomics methods have launched many international efforts, such as the Human Cell Atlas (*7*) initiative and the Tabula Sapiens (*8*) program. This has allowed studies to define differences in gene expression of same cell type, such as endothelial cells or immune cells, across different tissues and organs. A theme is that single cell types are shaped by tissue microenvironments (*82*) and thus allows studies on high level interaction networks to resolve the underlying tissue composition. Here, we show that the single cell data can complement and extend the information obtained from bulk tissue transcriptomics data published as part of the tissue section of the Human Protein Atlas (*13*). High similarity between the specificity classifications based on single cell and bulk tissue transcriptomics analysis demonstrates that bulk transcriptomics data gives an overall correct classification of most genes, but a more detailed and refined view is obtained when moving into single cell transcriptomics as done in this study.

Single cell transcriptomics has the potential to attribute gene expression specificity to a single cell type and subsets thereof, thus increasing the resolution at which gene expression specificity annotation can be performed, distinguishing cell type specific expression in unique single cell types present in only one organ from those that are identified across multiple organs. Furthermore, single cell transcriptomics has the potential to also identify expression in rare single cell types present at low levels, whose expression is not detected using bulk transcriptomics due to their low contribution to the total mRNA transcripts in the tissue sample. In summary, the comparison presented here between the single cell and the bulk tissue transcriptomics data show good agreement, but the cellular resolution obtained with the single cell data allows for insights of the same cell type across different tissues and thus accentuates the difference in single cell types in different tissues and organs.

Two new annotation tools are used in this analysis. First, the introduction of tau, representing the specificity of spatial expression across all analyzed single cell types as a continuous score, with a tau of 0 for a gene with equal expression across all single cell types and a tau of 1 for a gene exclusively expressed in one cell type. Second, the introduction of a new gene classification based on clustering of similar expression profiles across single cell types, which can be visualized in two dimensions using UMAP. These new tools are valuable contributions to earlier annotations based on bulk tissue transcriptomics (*13*), subcellular profiling (*83*), regional brain analysis (*37*), immune cell analysis (*36*), and blood protein analysis (*84*). A new open access version 21 of the Human Protein Atlas (www.proteinatlas.org) have been launched to enable researchers to explore the new genome-wide annotation on an individual gene level.

The cluster-based annotation of the human genome introduced here has the potential to increase the interpretability of the transcriptional landscape by labeling “neighborhoods” of similarly expressed genes by common features, in this case by specificity and function. This annotation is particularly consequential for uncharacterized genes, where hypotheses of functionality can be inferred from the transcriptional neighborhood. As an example, the cilia-related expression cluster described above harbors many genes with cilia-related function, but many genes lack functional annotation and these can be further validated for cilia function by the antibody-based profiling present in the tissue section of the Human Protein Atlas resource as exemplified by many “orf” genes (short for “open reading frame”) with no or little functional annotation.

In summary, we have added single cell data from 25 human tissues and integrated this data with different analytical platforms to create an open access resource to describe the expression of all human protein-coding genes across tissues and organs in the human body.

## Acknowledgements

We thank the entire staff of the Human Protein Atlas program and the Science for Life Laboratory (SciLifeLab) for their valuable contributions. This work was supported by the SciLifeLab & Wallenberg Data Driven Life Science Program (grant: KAW 2020.0239). The computations and data handling were enabled by resources provided by the Swedish National Infrastructure for Computing (SNIC) at UPPMAX partially funded by the Swedish Research Council through grant agreement no. 2018-05973. Main funding was provided from the Knut and Alice Wallenberg Foundation (WCPR).

## Author contributions

M.U., L.F. and C.L conceived and designed the analysis. M.K., M.B.A., M.S., C.Z., L.M., W.Z., R.S., F.H., A.D., B.K., J.V., E.S., M.B., F.E., A.M., J.M., L.B., Å.S., F.P., L.F., and C.L. collected and contributed data to the study. M.K., M.B.A., M.S., L.M., C.Z., L.F., A.M., performed the data analysis. K.v.F., M.Z., M.F., F.J. and P.O. created the database portal. M.U., L.F., C.L., F.P., Å.S., M.K., M.S., and M.B.A. drafted the manuscript. All authors read and approved the final manuscript.

## Supplementary data availability

All data needed to evaluate the conclusions in the paper are present in the paper and/or the Supplementary Materials. Single cell and bulk tissue gene expression data are available to download on the Human Protein Atlas resource download page (https://proteinatlas.org/about/download).

## Methods

### Single cell transcriptomics datasets

The single cell RNA sequencing dataset is based on meta-analysis of literature on single cell RNA sequencing and single cell databases that include healthy human tissue. To avoid technical bias and to ensure that the single cell dataset can best represent the corresponding tissue, the following data selection criteria were applied: (1) Single cell transcriptomic datasets were limited to those based on the Chromium single cell gene expression platform from 10X Genomics (version 2 or 3); (2) Single cell RNA sequencing was performed on single cell suspension from tissues without pre-enrichment of cell types; (3) Only studies with > 4,000 cells and 20 million read counts were included, (4) Only dataset whose pseudo-bulk transcriptomic expression profile is highly correlated with the transcriptomic expression profile of the corresponding HPA tissue bulk sample were included. It should be noted that exceptions were made for lung (∼7.3 million reads), pancreas (3,719 cells) and rectum (3,898 cells) to include various single cell types in the analysis.

In total, single cell transcriptomics data for 25 tissues and peripheral blood mononuclear cells (PBMCs) were analyzed. These datasets were respectively retrieved from the Single Cell Expression Atlas (https://www.ebi.ac.uk/gxa/sc/home), the Human Cell Atlas (https://www.humancellatlas.org), the Gene Expression Omnibus (https://www.ncbi.nlm.nih.gov/geo/), the Allen Brain Map (https://portal.brain-map.org/), and the European Genome-phenome Archive (https://www.ebi.ac.uk/ega/). The complete list of references is shown in **Supplementary data S1**.

### Clustering of single cell transcriptomics data

For each of the single cell transcriptomics datasets, the quantified raw sequencing data were downloaded from the corresponding depository database based on the accession number provided by the corresponding study in the available format (total cells, read, and feature counts, or count tables). Unfiltered data were used as input for downstream analysis with an in-house pipeline using Scanpy (version 1.4.4.post1) in Python 3.7.3 for the 13 tissues and PBMC published in HPA v20 and Scanpy (version 1.7.2) in Python 3.8.5 for the 12 tissues published in HPA v21. In the pipeline, the data were filtered using two criteria: a cell is considered as valid if at least 200 genes are detected and a gene is considered as valid if it is expressed in at least 10% of the cells. Specially, in the HPA v21, tissues which contain more than 10,000 cells used 1000 cells as their cutoff. Subsequently, the cell counts were normalized to have a total count per cell of 10,000. The valid cells were then clustered using Louvain clustering function within Single-Cell Analysis in Python (Scanpy). Default values of parameters were used in clustering. More in detail, the features of cells were projected into a PCA space with 50 components using UMAP, and a k-nearest neighbours (KNN) graph was generated. 15 neighbours were used in the network for Louvain, while the resolution of clustering was set as 1.0. The total read counts for all genes in each cluster was calculated by adding up the read counts of each gene in all cells belonging to the corresponding cluster. Finally, the read counts were normalized to transcripts per million protein coding genes (pTPM) for each of the single cell clusters. When calculating the expression profile for pseudo-bulk samples based on single cell transcriptomics, we added the read counts for all genes from all cells of the sample, and normalized it to pTPM in the same way as for the cluster ones.

### Single cell type annotation

Each of the 444 different cell type clusters was manually annotated based on an extensive survey of well-known tissue- and cell type–specific markers, including both markers from the original publications and additional markers used in pathology diagnostics. For most single cell types, three marker genes were used, and for each cluster, one main cell type was chosen based on the overall expression pattern of all the marker genes. For some clusters, no main cell type could be selected, and these clusters were not used for classification. The most relevant markers (**Supplementary data S2**) are presented in a heatmap on the in the Single Cell Type section on each organ- and gene-specific page to clarify cluster annotation to visitors.

### Tissue transcriptomics datasets

Non-brain human tissue samples (n = 177) for analysis of bulk RNA-seq and protein expression in the HPA datasets were collected and handled in accordance with Swedish laws and regulation. Tissues were obtained from the Clinical Pathology department, Uppsala University Hospital, Sweden and collected within the Uppsala Biobank organization. All samples were anonymized for personal identity by following the approval and advisory report from the Uppsala Ethical Review Board (Ref # 2002-577, 2005-388, 2007-159, 2011-473). The RNA extraction and RNA-seq procedure has been described previously (*13*). For brain samples (n=1324), samples covering 206 regions of the brain were collected using the punch needle approach (*39*). After RNA extraction and ribosomal RNA depletion RNA was quantified followed by a normalization step to correct for donor and technical variation. All brain samples were collected by the Human Brain Tissue Bank (HBTB; Semmelweis University, Budapest) and the Human Brain Research Laboratory (ELKH Institute of Experimental Medicine). The activity of the HBTB has been authorized by the Committee of Science and Research Ethic of the Ministry of Health Hungary (ETT TUKEB: 189/KO/02.6008/2002/ETT) and the Semmelweis University Regional Committee of Science and Research Ethic (No. 32/1992/TUKEB) to remove human brain tissue samples, collect, store and use them for research. Additionally, RNA-seq data was collected from GTEx (n = 10,442), and all data were summarized as TPM by mapping processed reads to the human reference genome GRCh38.p13 and with gene models based on Ensembl (v103) (*85*) using Kallisto (v.0.46.2) (*86*). Next, the gene expression levels were calculated by summing up all the TPM values of all alternatively spliced protein coding transcripts of the corresponding gene for a total number of 20,090 protein-coding genes and rescaled to sum to one million, forming protein-coding transcripts per million (pTPM), and subsequently normalized using Trimmed Means of M (TMM) using NOISeq’s (*87*) tmm function with a median column as reference, with the parameters logratioTrim = 0.3, sumTrim = 0.3, doWeighting = F, to generate normalized transcripts per million (nTPM).

### Prediction of secreted and membrane location

The results of a whole-proteome analysis using seven different prediction algorithms for membrane topology and three methods for signal peptide (SP) prediction was used to create majority decision-based methods to estimate the human secreted and membrane proteomes (*88*). For each gene, the splice variants with at least one transmembrane (TM) segment with overlapping predictions by four out of the seven methods were considered membrane proteins. Splice variants with a predicted SP by at least two of the three methods, not predicted as membrane protein according to above, as well as not predicted to reside in intracellular locations, such as Golgi or Endoplasmic reticulum, according to our previous manual annotation (*84*), were considered secreted proteins. Splice variants not classified according to above criteria were considered intracellular proteins.

The predictions for the splice variants were then used to classify the genes into four different predicted location categories. Genes with only secreted or a combination of secreted and intracellular splice variants were classified as ‘secreted’, genes with only membrane or a combination of membrane and intracellular splice variants were classified as ‘membrane’, genes with both secreted and membrane splice variants, as well as genes having all three types of splice variants were classified as ‘membrane/secreted’, and genes with only intracellular splice variants were classified as ‘intracellular’.

### Antibody based protein profiling

Immunohistochemical staining and high-resolution digitization of stained tissue microarray (TMA) slides were performed essentially as previously described (*89*). Primary antibodies were diluted and optimized based on IWGAV (International Working Group for Antibody Validation) criteria for antibody validation (*90*). Antibodies used for the immunohistochemical example images are listed in **Supplementary data S6**. Protocol optimization was performed on a test TMA containing 20 different normal tissues. The stained slides were digitized with Scanscope AT2 (Leica Aperio, Vista, CA, USA). All images were manually evaluated by two independent observers, comprising a total of 576 images per antibody, covering 15,323 genes available on v21.proteinatlas.org.

### Gene specificity classification

The expression data from single cell and bulk was aggregated from 444 single cell type clusters into 76 unique single cell types and from 55 tissue types into 36 consensus tissues, representing the main single cell types and tissues in the whole human body. For both datasets, the aggregated expression data (nTPM) was used to classify each gene into one of the following categories: (i) cell type/tissue enriched, (ii) group enriched, (iii) cell type/tissue enhanced, (iv) low cell type/tissue specificity and (v) not detected, according to the criteria in Supplementary table 1. Additionally, log10(nTPM + 1) transformed values were used to calculate the “tau” specificity score (*43*).

### Gene expression clustering

In short, gene expression clustering was performed for the single cell and tissue data independently. First, expressed genes (nTPM >1) were selected (n = 18,965 in single cell and n= 19,023 in tissue) and scaled to z-scores to account for differences in dynamic range across single cell types. The data was normalized, and principal component analysis (PCA) was used to reduce the dimensionality of the data for noise reduction, selecting the components which retain most of the variation of gene expression across samples. The lower-dimensional data was used to calculate the Spearman distance between genes (1 – Spearman correlation), which were then transformed into a shared nearest neighbour (SNN) graph using the FindNeighbors function in Seurat, with the k.param = 20 and the prune.SNN parameter set to 1/15 for tissue and 1/20 for single cell. Nearest neighbors were subsequently visualized as a UMAP plot.

Alternative gene clusterings were performed for the single cell and tissue datasets using the Louvain clustering implementation in Seurat, using the FindClusters function with the resolution parameter ranging from 0.5 to 20 in order to explore a wide range of alternatives for the number of clusters. Additionally, each of the resolutions was run 100 times to account for the stochasticity of the clustering process, and the results were then integrated into a consensus clustering using the cl_consensus function in the clue package with the parameter method = “SE”. The final resolution was selected according to the stability of the 100 alternative clustering generated calculated using the adjusted rand index (ARI).

Analysis of cluster-associated specificity and function was done by performing one-sided hypergeometric tests (overrepresentation analysis) towards terms within Gene Ontology (GO) Biological Process (BP), Cellular Component (CC), and Molecular Function (MF) (*45, 46*); pathways in Reactome (*91*) and KEGG (*92–94*); single cell type markers from PanglaoDB (*44*); TRRUST transcription factors (*95*); and annotation made in the Human Protein Atlas, including specificity classifications across tissues (*13*), single cell types (*14*); subcellular protein location assessed with immunofluorescence in cell lines (*83*); predicted protein class (intracellular, secreted, and membrane) based on signaling peptides (*13*); and secretome locations (tissues and body liquids to which secreted proteins are targeted, such as blood) (*84*). This overrepresentation analysis generated statistically significant gene overlaps between each cluster and the beforementioned gene annotations, which gives a comprehensive picture of shared functions of cluster genes. The overrepresentation results, together with the lists of genes and their expression profiles for each of the clusters were carefully inspected by experts which assigned a main specificity and function associated to a reliability score indicating the confidence of the assignment.

In order to visualize the expression clusters, the SNN graph was used as an input for dimensionality reduction using UMAP. The areas comprising most of the cluster genes are summarised using polygons to facilitate the visualization of clusters.

### Data visualization

Data visualization was performed in R (v.4.0.3) (*96*), using the ggplot2 (v 3.3.5) (*97*), and the complementary packages concaveman (v 1.1.0), dendextend (v 1.15.1) (*98*), ggalluvial (v 0.12.3) (*99*), ggbeeswarm (v 0.6.0), ggdendro (v 0.1.22), scatterpie (v 0.1.7), ggraph (v 2.0.5), ggrastr (v 0.2.3), ggrepel (v 0.9.1), ggupset (v 0.3.0), patchwork (v 1.1.1), pcaMethods (v 1.82.0) (*100*), rrvgo (v 1.2.0), tidygraph (v 1.2.0), treemapify (v 2.5.5), uwot (v 0.1.10), and viridis (v 0.6.1). The figures were assembled in Affinity designer (v 1.10.0.1127).

## Supplementary Materials for

**Figure S1.**
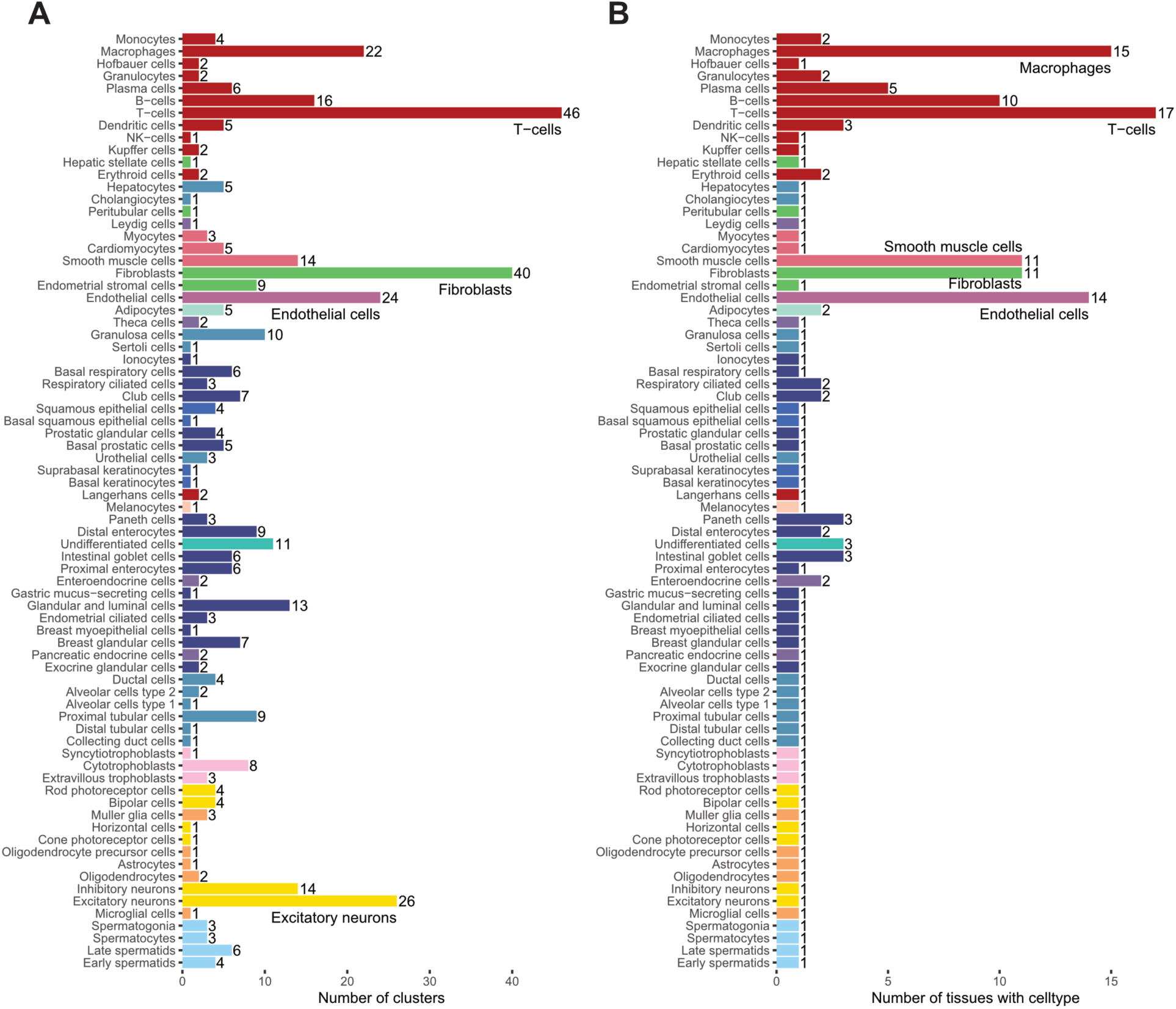
Pooling of single cell type clusters. **A)** Barplot showing the number of single cell type expressions clusters (n = 444) pooled into unique single cell types (n = 76). **B)** Barplot showing the number of tissues where the single cell types are represented.

**Figure S2.**
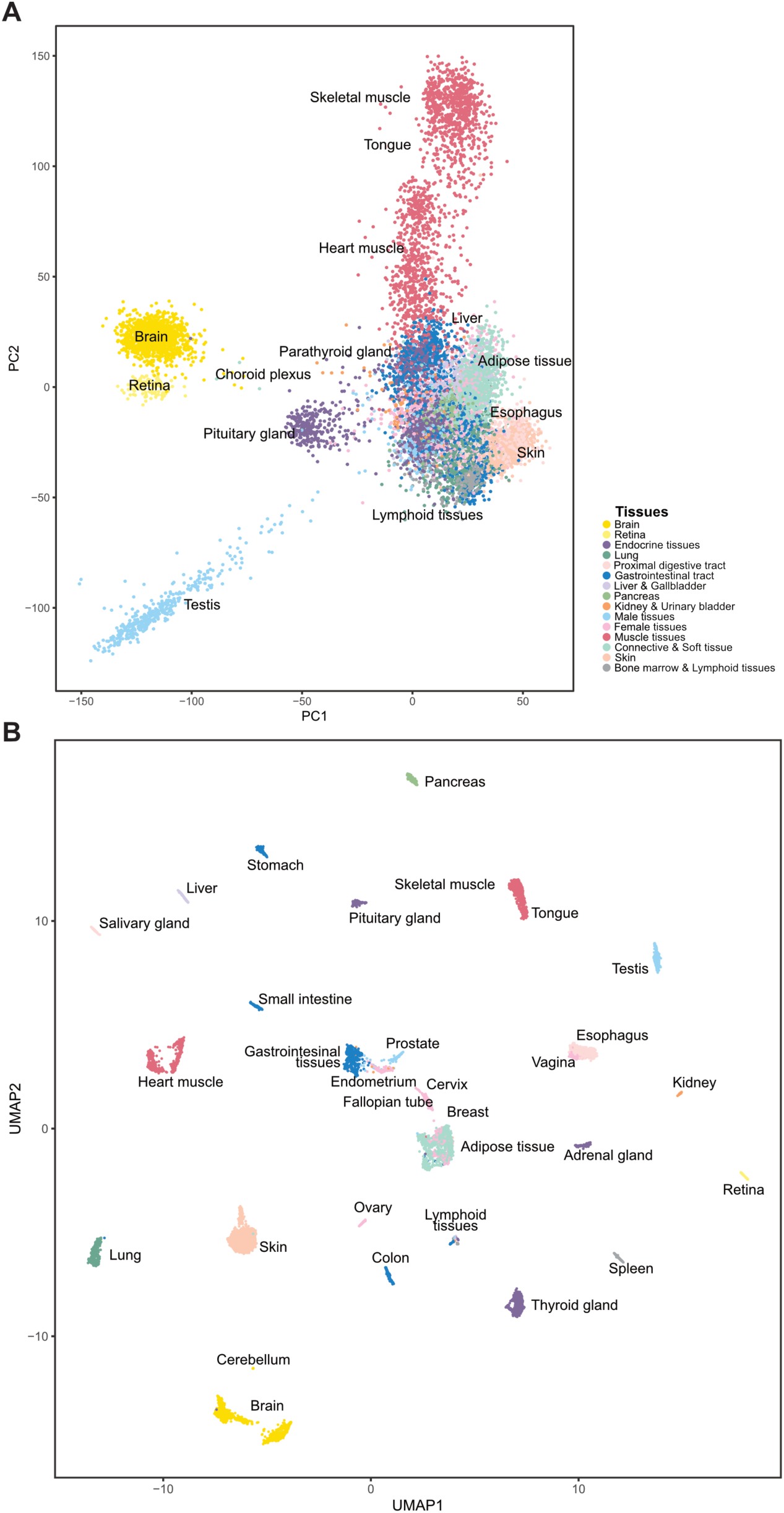
Bulk transcriptomics data. (A) PCA and (B) UMAP of 11,794 tissue samples. Points are colored by tissue type.

**Figure S3.**
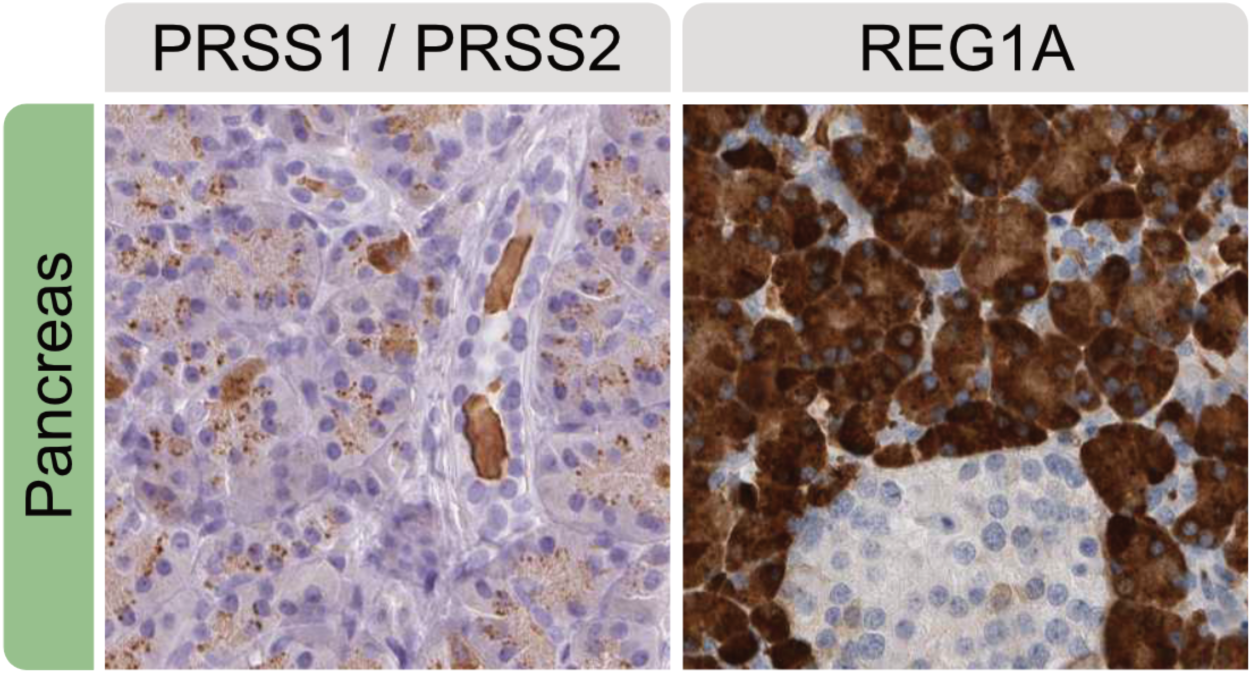
IHC of PRSS1, PRSS2, and REG1A in pancreas.

**Figure S4.**
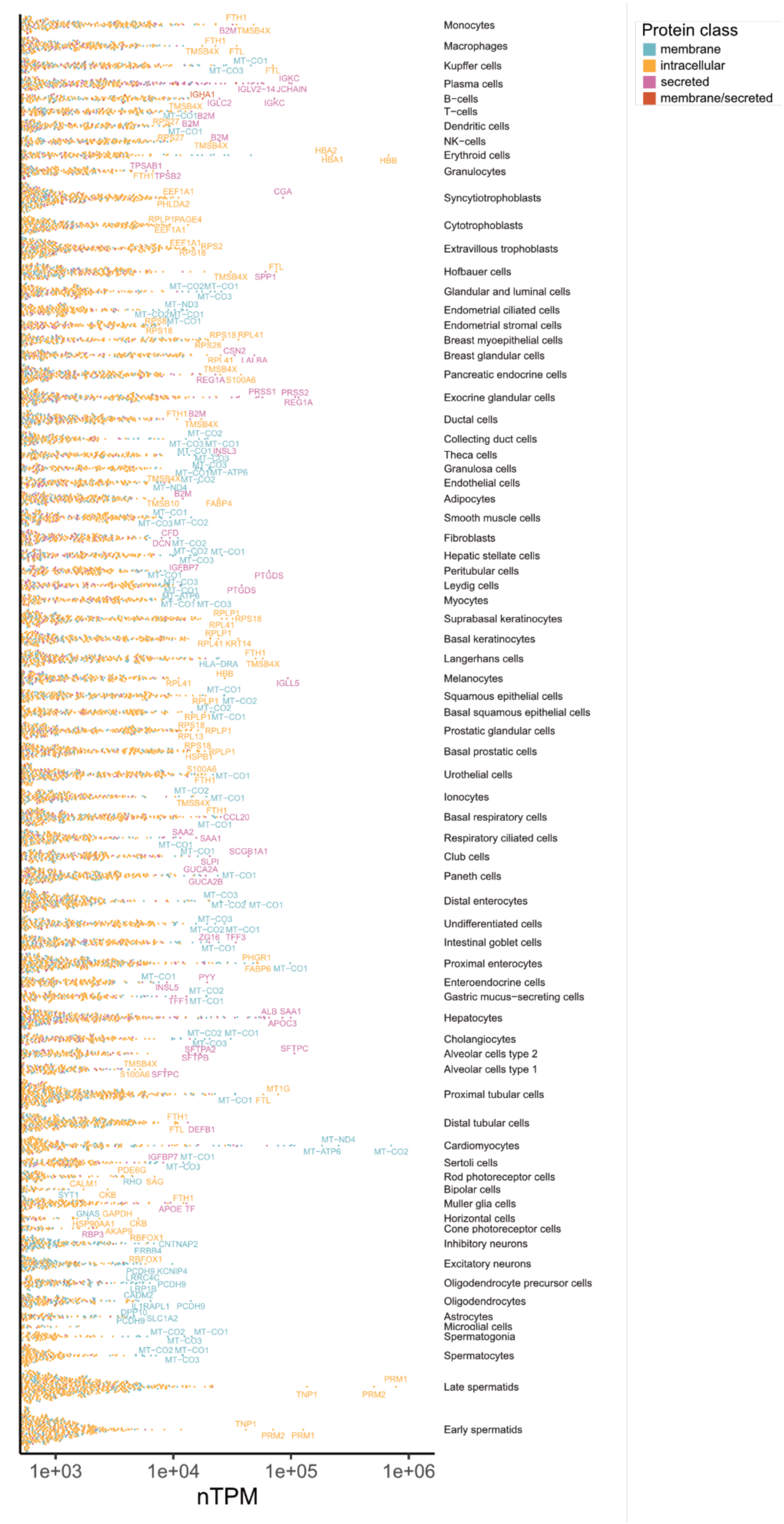
Expression levels of genes expressed > 500 nTPM across all 76 cell types. The top three expressed genes are labeled and all genes are colored according to the protein class based on prediction of membrane, intracellular, secreted or membrane/secreted proteins.

**Figure S5.**
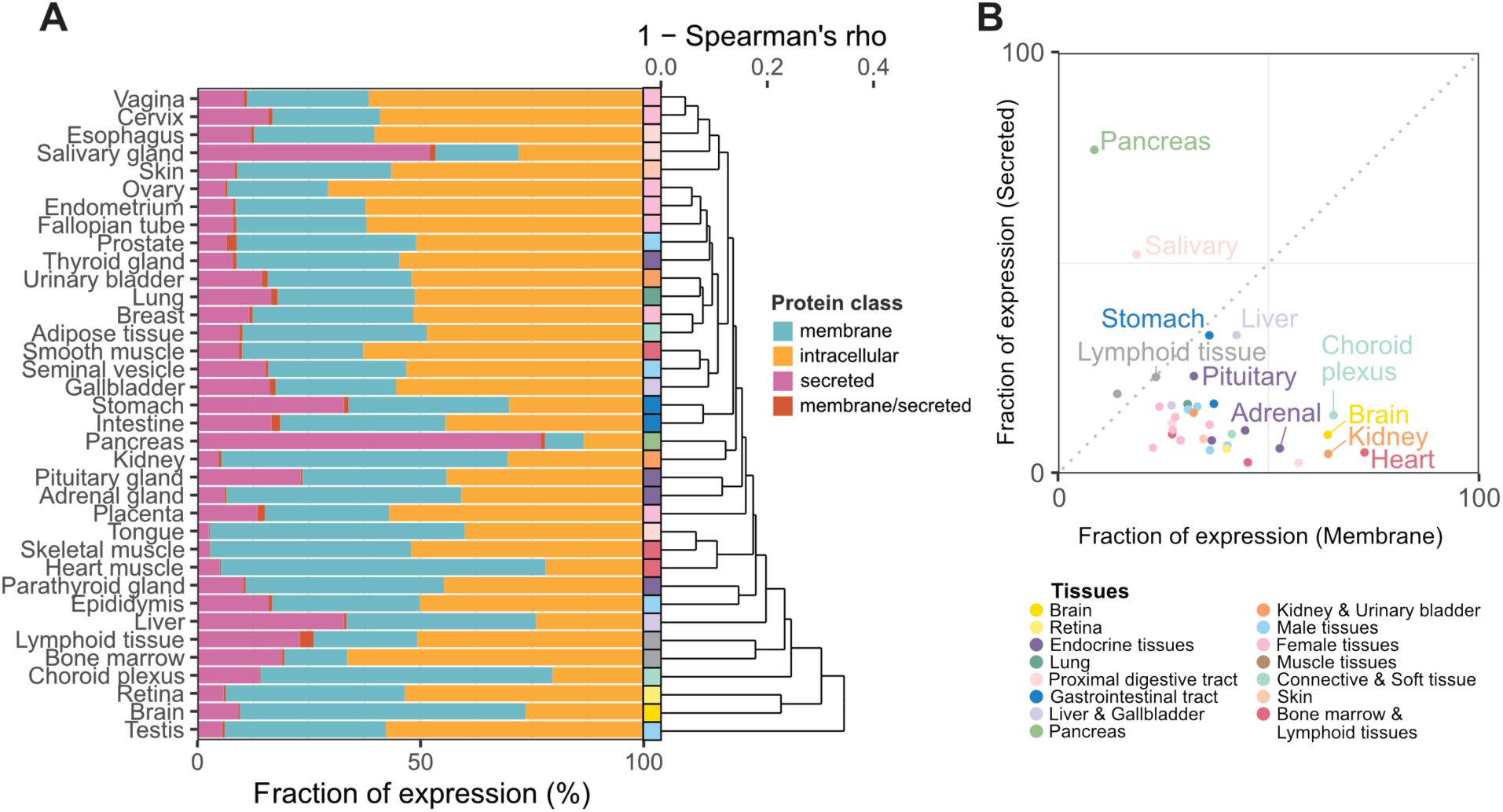
Fraction of transcripts corresponding to protein classes in tissues from bulk transcriptomics. **(A)** Barplot showing the fraction of total gene expression in each tissue type corresponding to genes predicted to encode membrane, intracellular, secreted or membrane/secreted proteins, respectively. The dendrogram is based on the average distance (1 – Spearman’s rho) between the 36 tissue types. **(B)** Relationship between the fraction of total gene expression corresponding to secreted and membrane protein encoding genes in the 36 tissue types.

**Figure S6.**
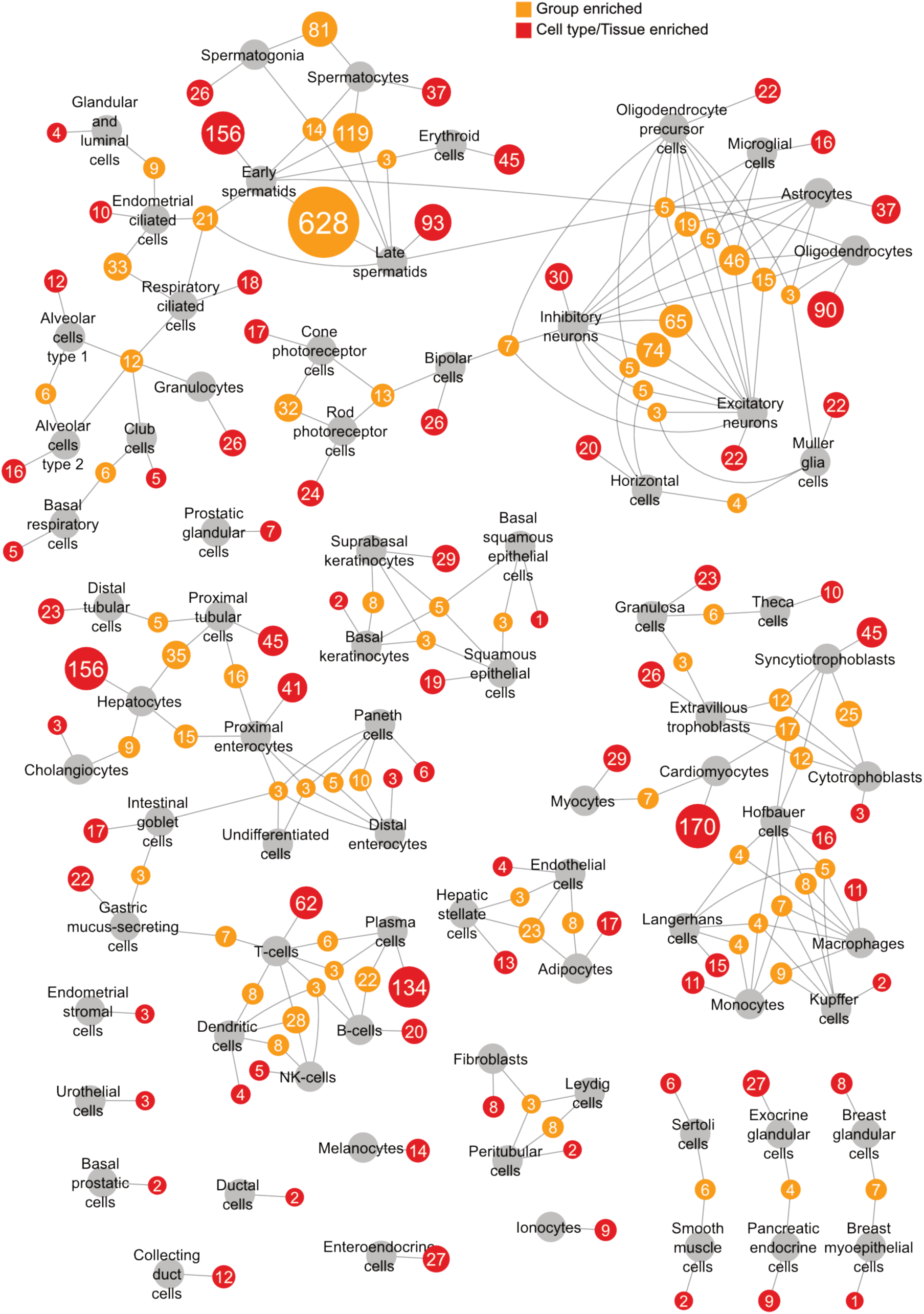
Network showing the number of enriched genes across cell types. Cell types are connected with nodes indicated the number of cell type enriched genes (red), and group enriched genes (yellow) per each cell type. For group enriched genes, the connected cell types indicate which combination of cell types these genes are co-enriched in. Nodes are filtered to reduce complexity: All nodes of cell type enriched genes are shown, and nodes that contain among the top two highest number of genes for any connected cell type and contains at least 3 genes.

**Figure S7.**
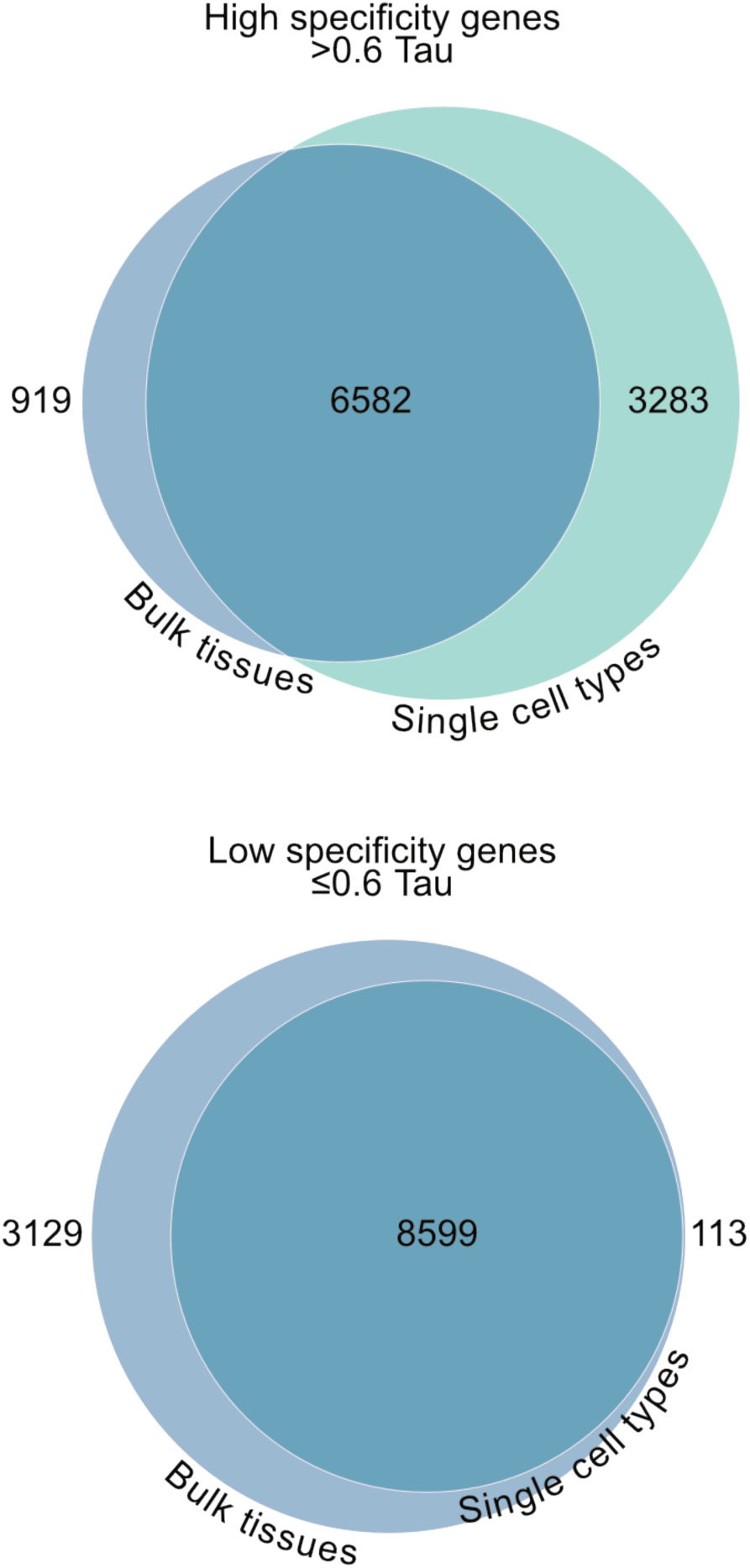
Overlap between the high and low specificity genes between single cells and bulk tissues based on tau. Venn diagrams showing the overlap of genes with high specificity (top) and low specificity (bottom) between single cell and bulk tissue transcriptomics, using a cutoff of 0.6 tau.

**Figure S8.**
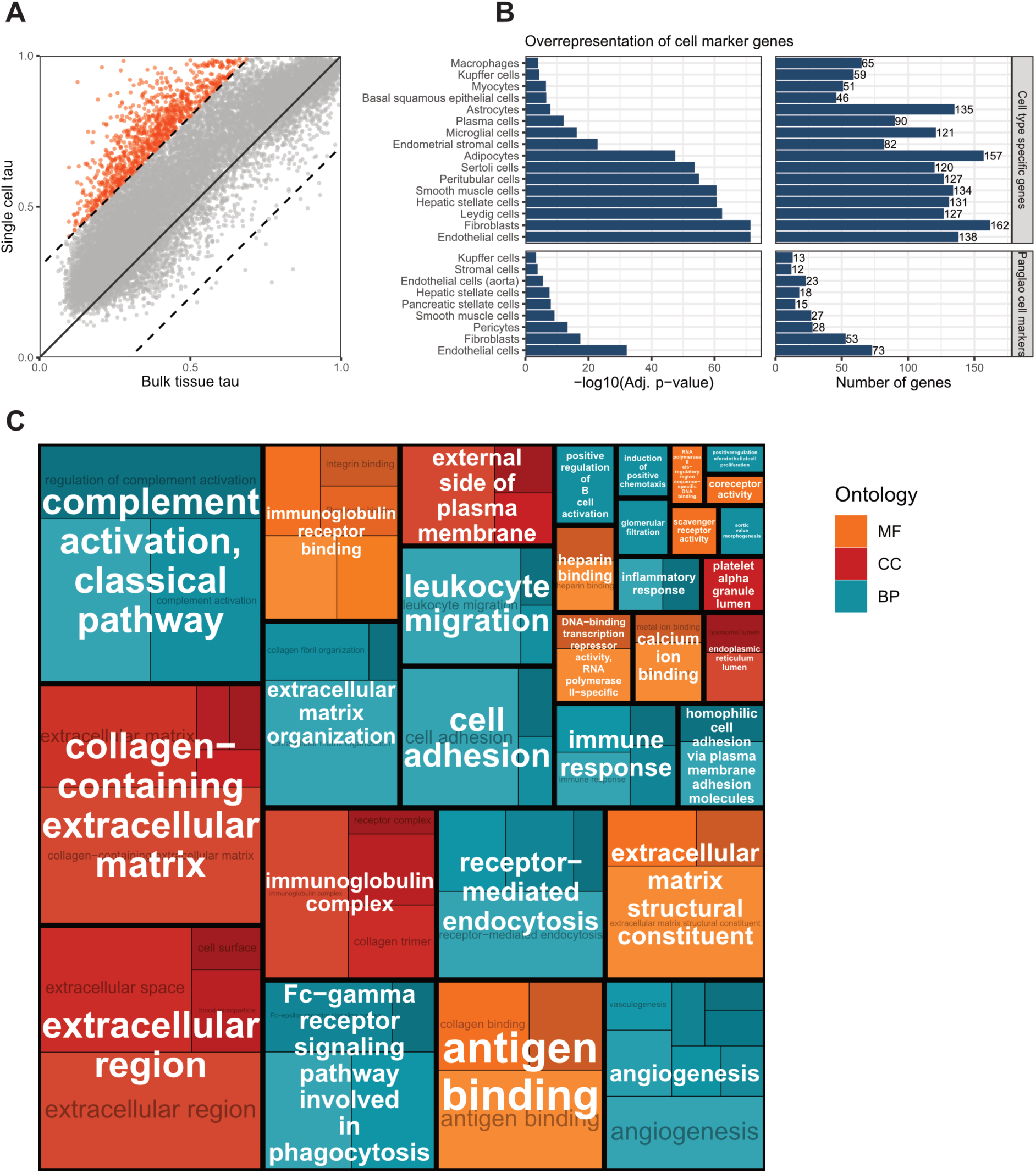
Overrepresentation analysis of genes with higher specificity in single cell types compared to tissues. **(A)** Comparison of tau scores in single cell and bulk transcriptomics datasets. Genes with higher specificity (>0.3 higher tau) in single cell compared to bulk tissues are colored in red (n = 1073). **(B)** Gene set enrichment analysis showing the overrepresentation of cell markers among the 1073 genes with higher specificity across single cells compared to across bulk tissues, showing Benjamini-Hochberg (BH) adjusted p-value (left) and number of genes (right) for cell types with statistically significant (adj. p < 0.001) overrepresentation of cell specific markers according to internal (HPA, top) and external (PanglaoDB, bottom) specificity annotations. **(C)** Treemap of GO gene set enrichment analysis of the 1073 genes with higher specificity across cell types compared to bulk tissue types. The box size for each GO term is proportional to the log-transformed BH adjusted p-values for significantly enriched terms (adj. p < 0.05), and terms with substantial gene overlap are grouped to parent terms. Boxes are colored by the three GO domains: MF = Molecular function, CC = Cellular component, BP = Biological process.

**Figure S9.**
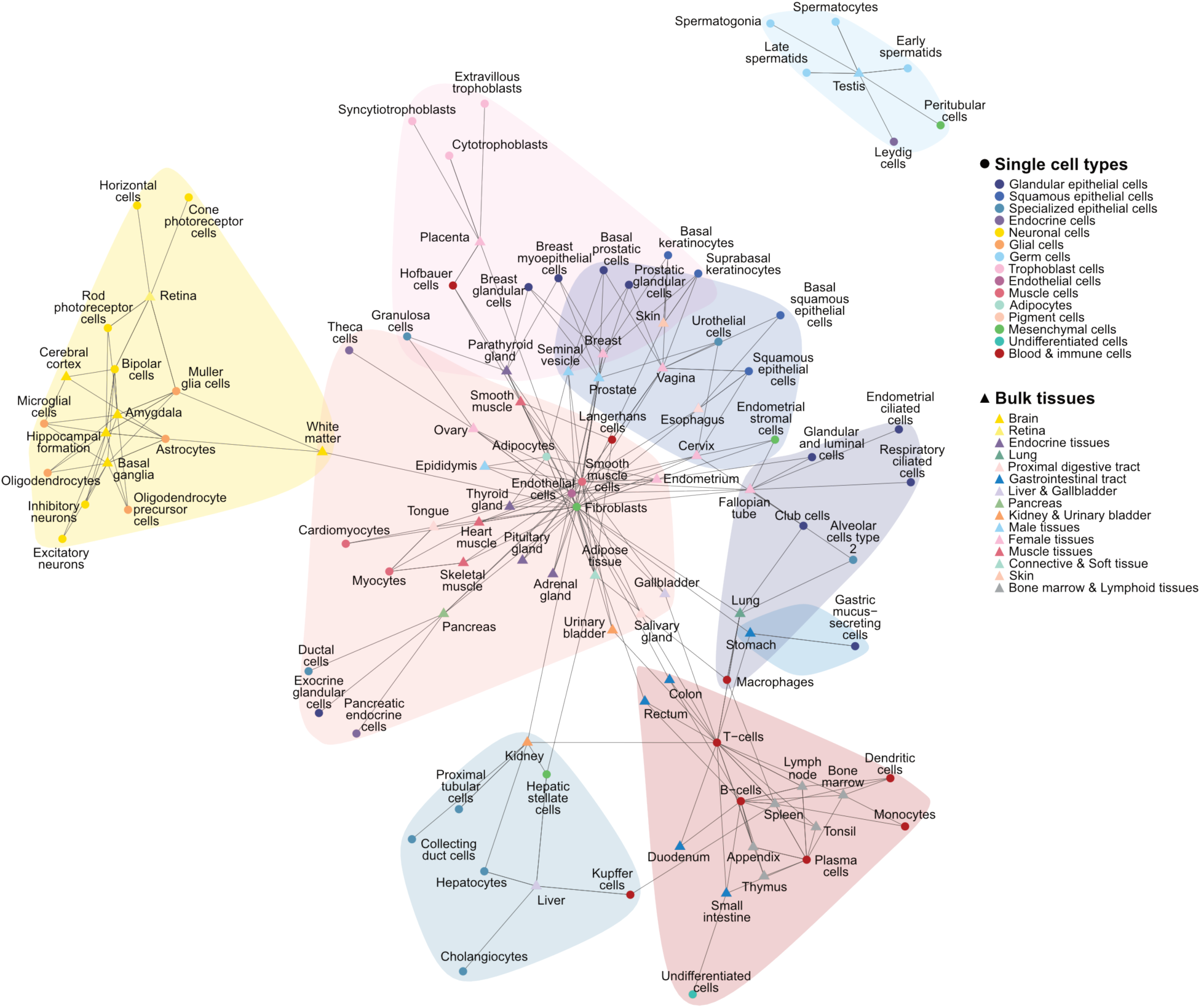
Correlation network showing the pairwise most similar cell types and tissues. Correlation calculated between single cell types and bulk tissues for genes with enriched expression in at least one single cell type or tissue. Top three Spearman correlations for each node (with a cutoff of correlation >0.6) are shown.

**Figure S10.**
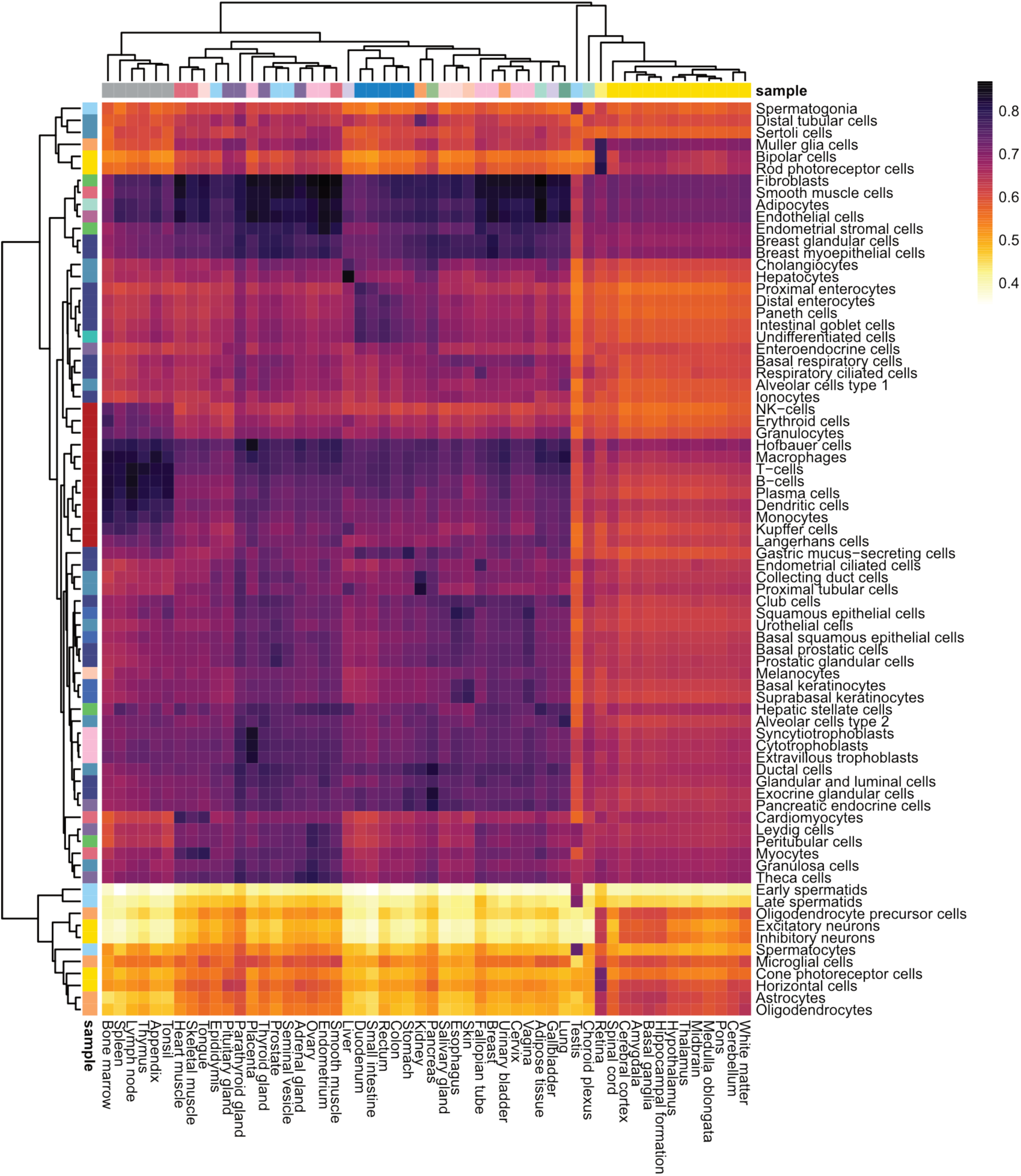
Spearman correlation heatmap between single cell types (y-axis) and bulk tissues (x-axis). Spearman correlation is calculated across all genes. Dendrograms result from hierarchical clustering using Ward’s criterion.

**Figure S11.**
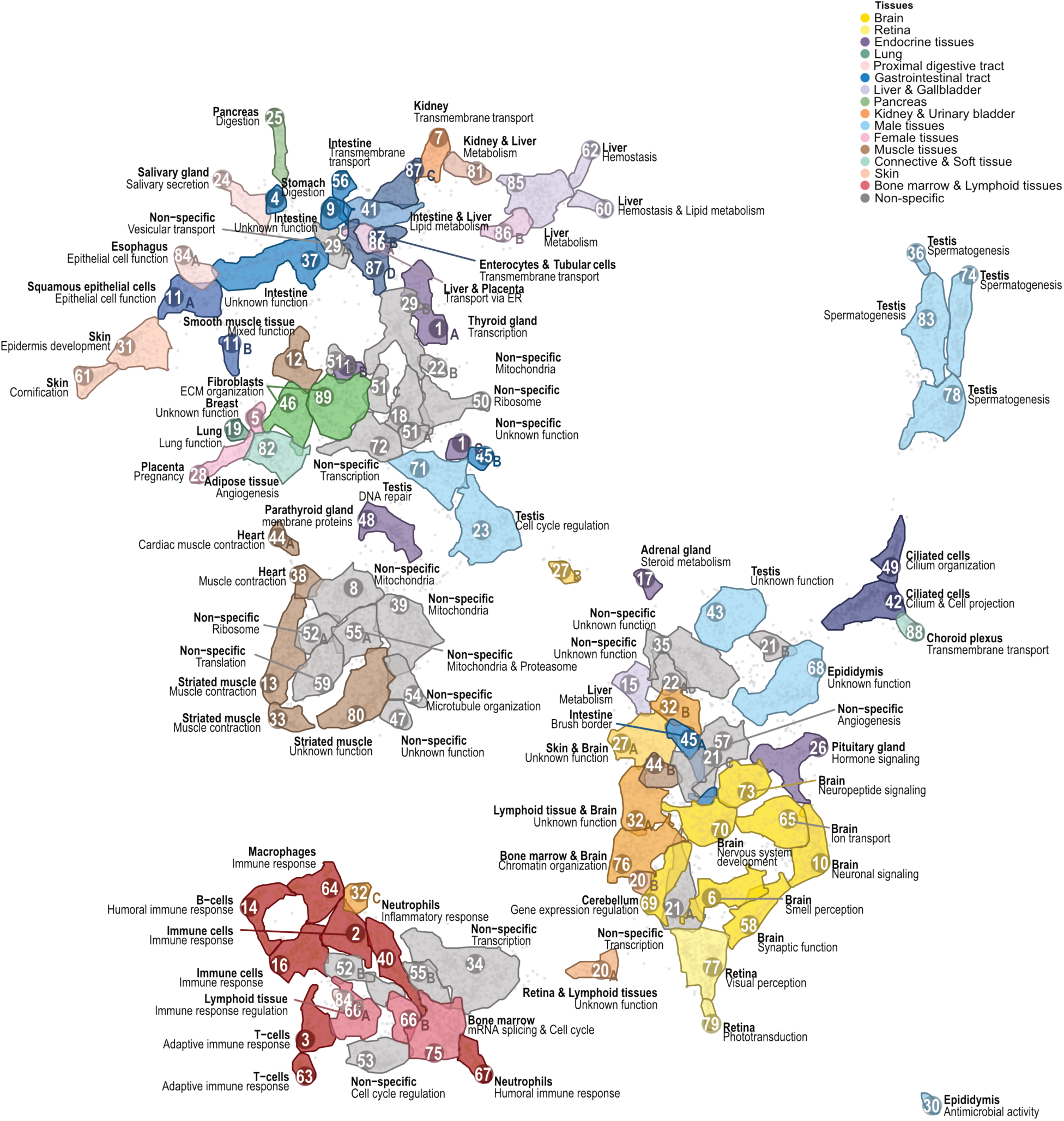
UMAP visualization of the bulk tissue expression clusters. Clusters are summarized as polygons showing the areas in the UMAP where most genes of the cluster are located. Cluster polygons are colored according to the bulk tissue type specificity that the cluster’s genes exhibit, while non-specific clusters are colored grey, and are labelled according to the annotated specificity (bold) and function.

**Figure S12.**
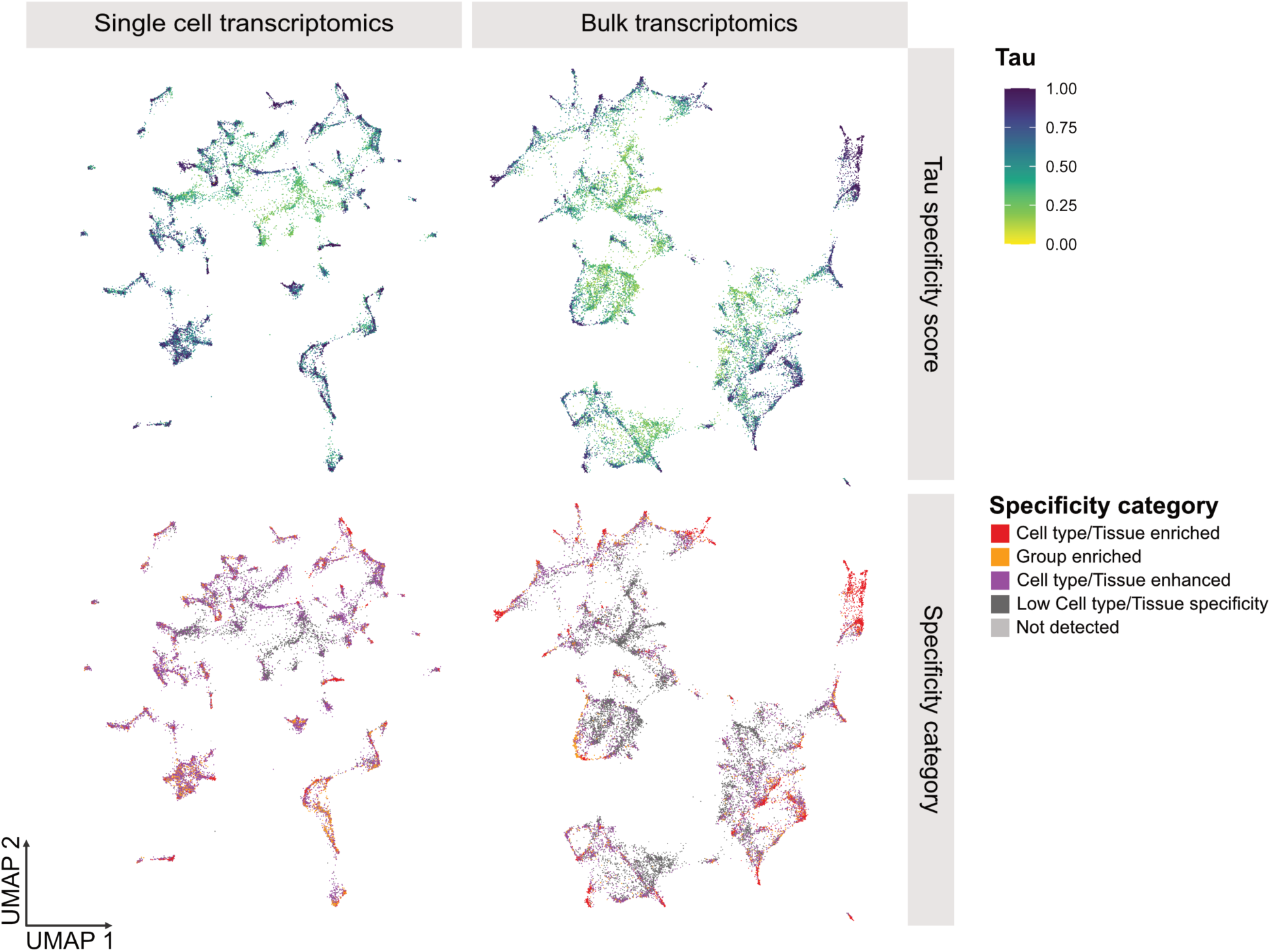
Tau score and specificity category superimposed on single cell and bulk tissues gene-based UMAP results. Each point corresponds to a gene and is colored by its specificity according to tau specificity score (top) and specificity classification (bottom).

**Figure S13.**
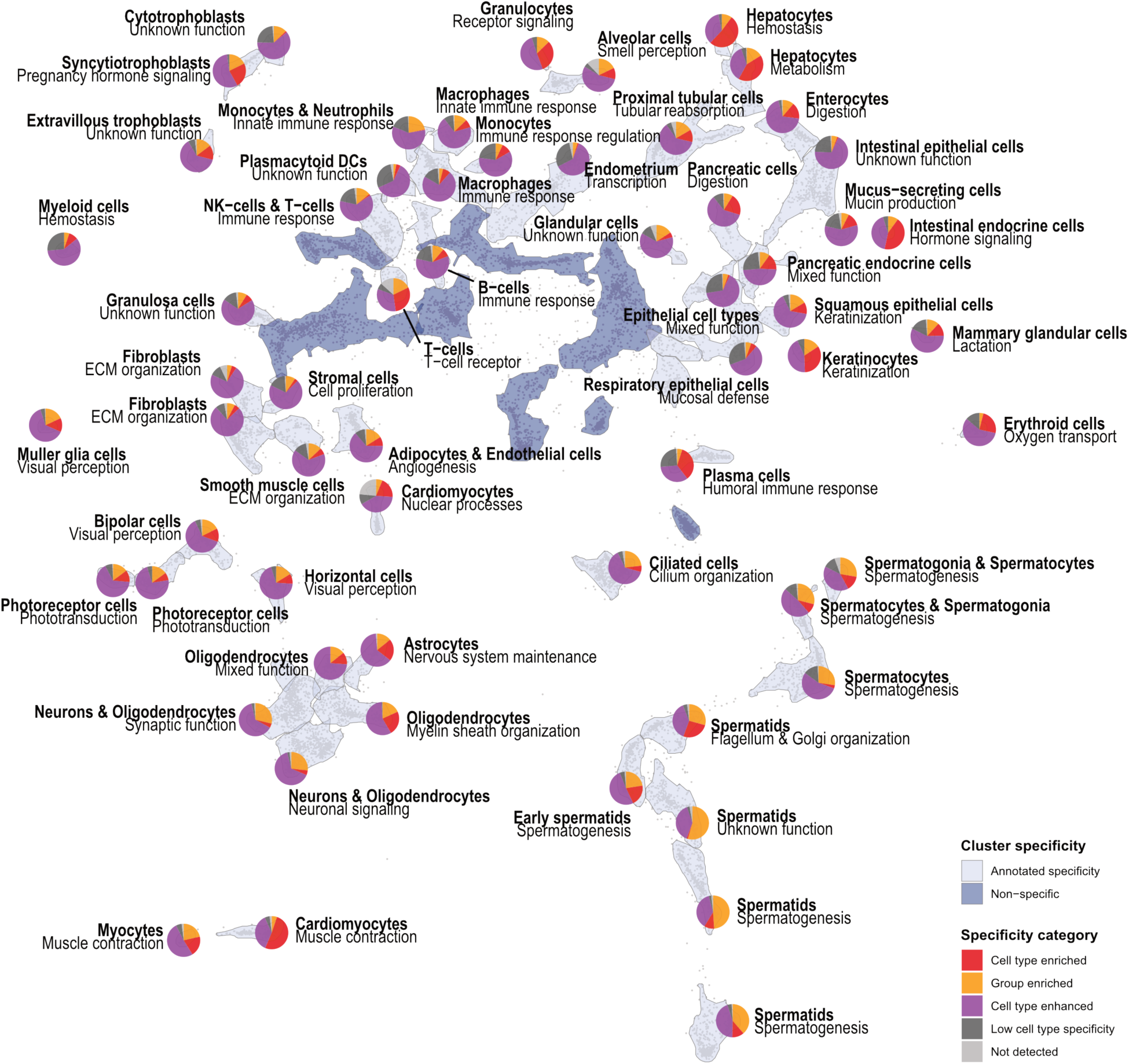
Scatterpie plot showing the proportion of genes from each specificity category in cell type expression clusters with annotated specificity. Each pie shows the fraction of all genes in the cluster in each of the five specificity categories.

**Figure S14.**
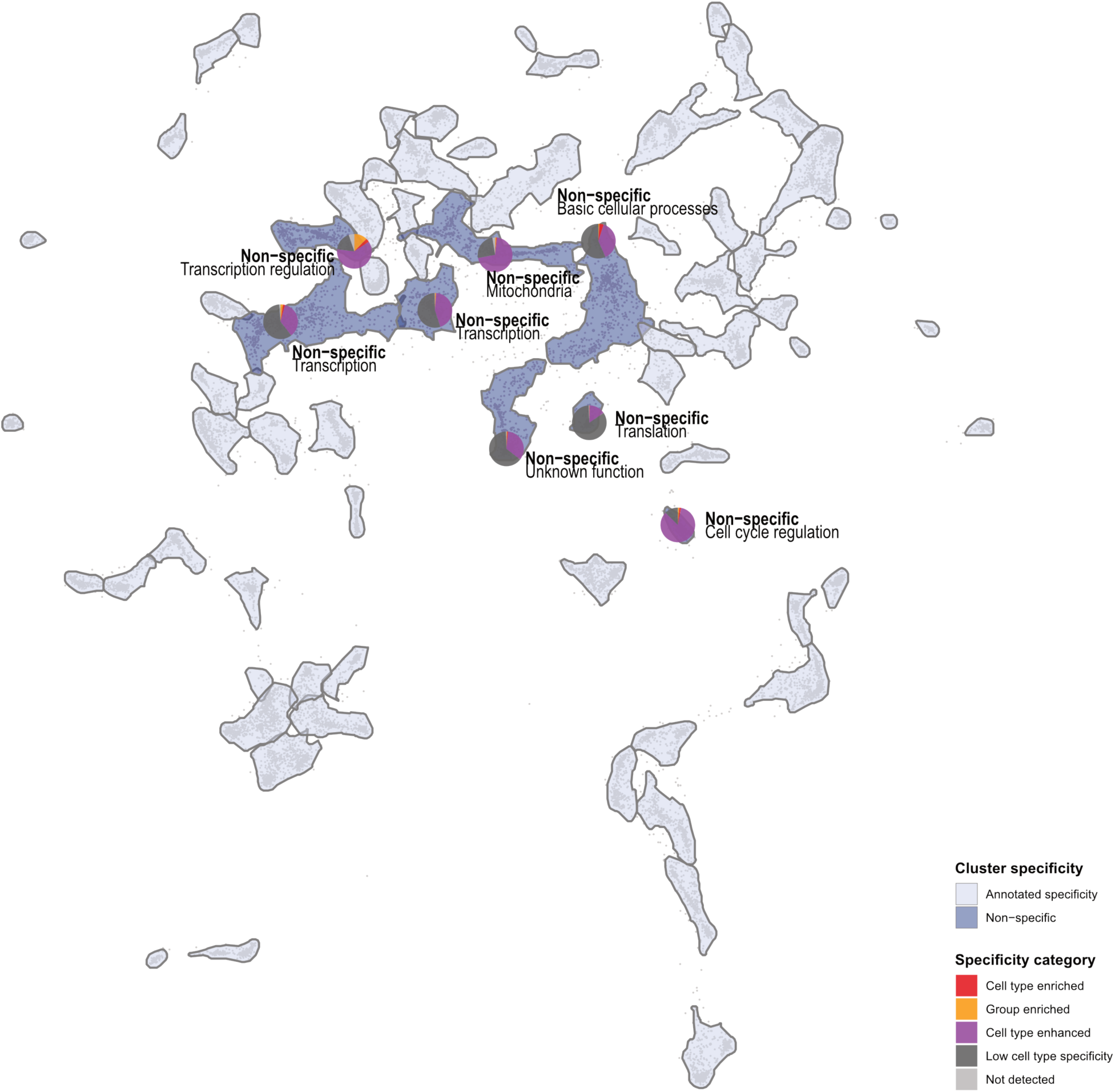
Scatterpie plot showing the proportion of genes from each specificity category in cell type expression clusters annotated as non-specific. Each pie shows the fraction of all genes in the cluster in each of the five specificity categories.

**Figure S15.**
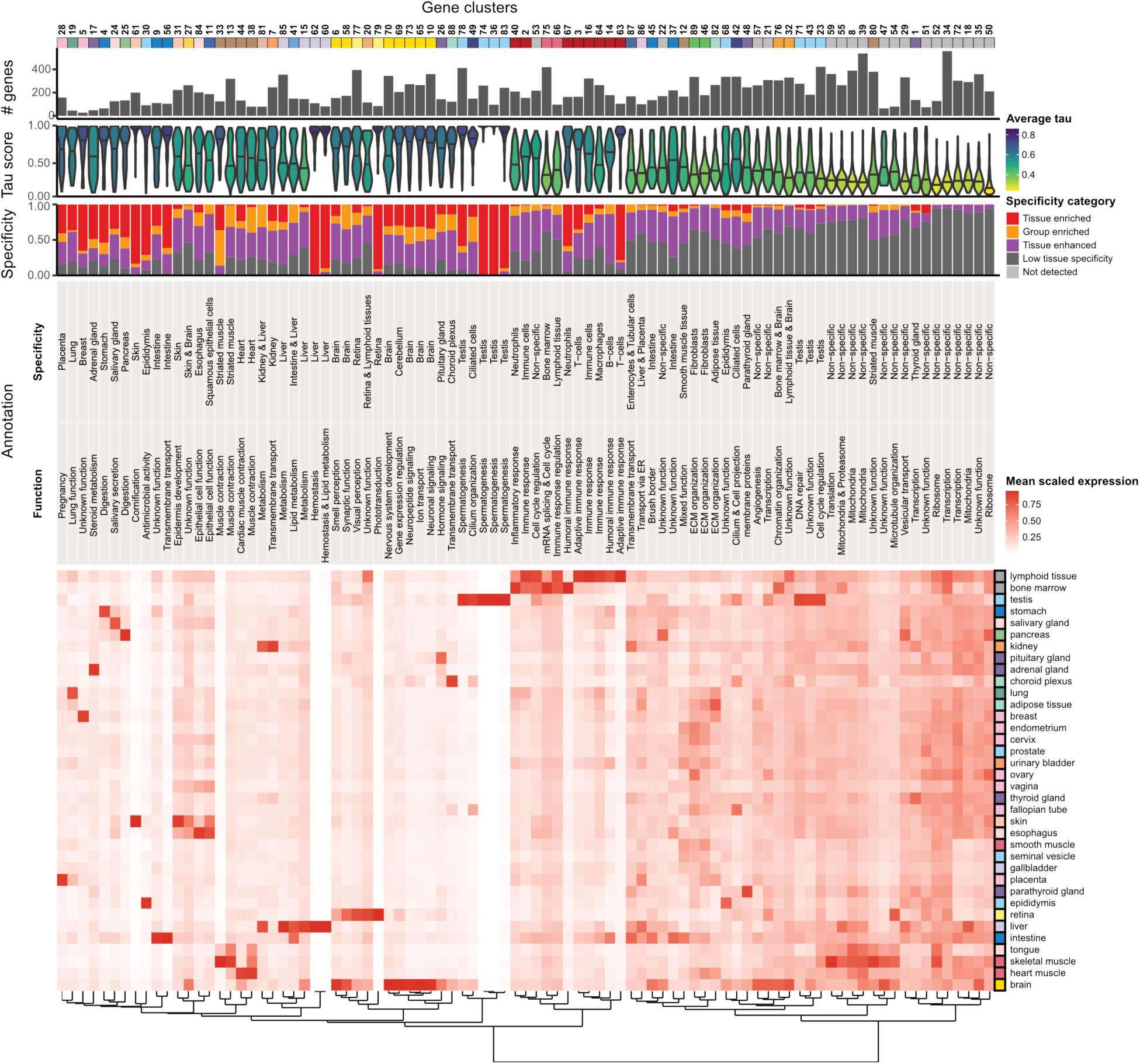
Characterization of bulk tissue expression clusters. The number of genes, distribution of tau scores, proportion of genes in each specificity category, annotated specificity and function and summarized mean expression across the 36 consensus tissues is indicated for each of the 89 clusters. The dendrogram is obtained by hierarchical clustering of distances based on gene expression levels for all tissues using Ward’s criterion.

**Figure S16.**
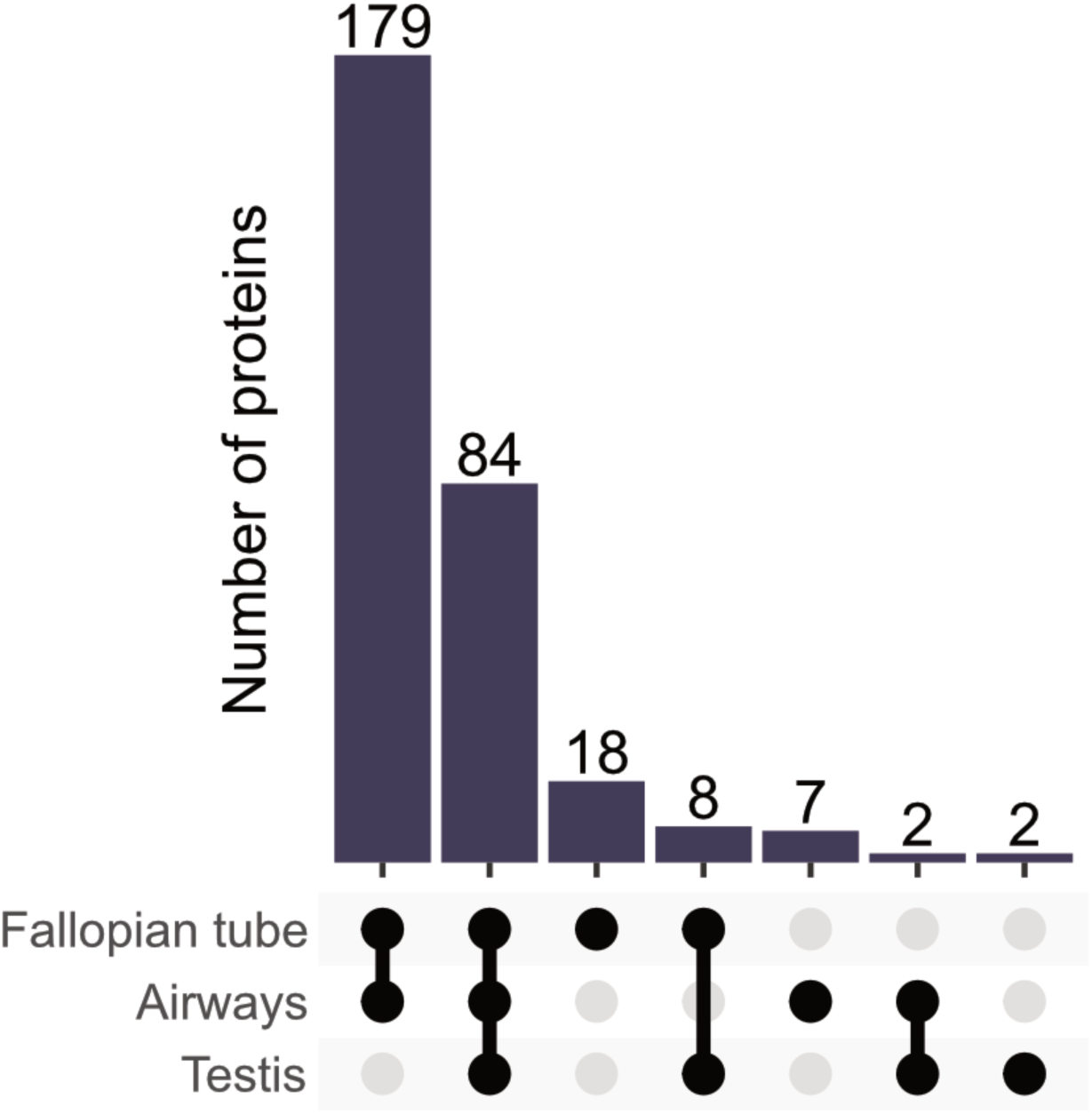
The number of proteins from the cilia related gene expression cluster 49 with staining in cilia across tissues. Upset barplot showing the number of proteins with staining in ciliated cell in combinations of tissues as indicated by the x-axis.

**Figure S17.**
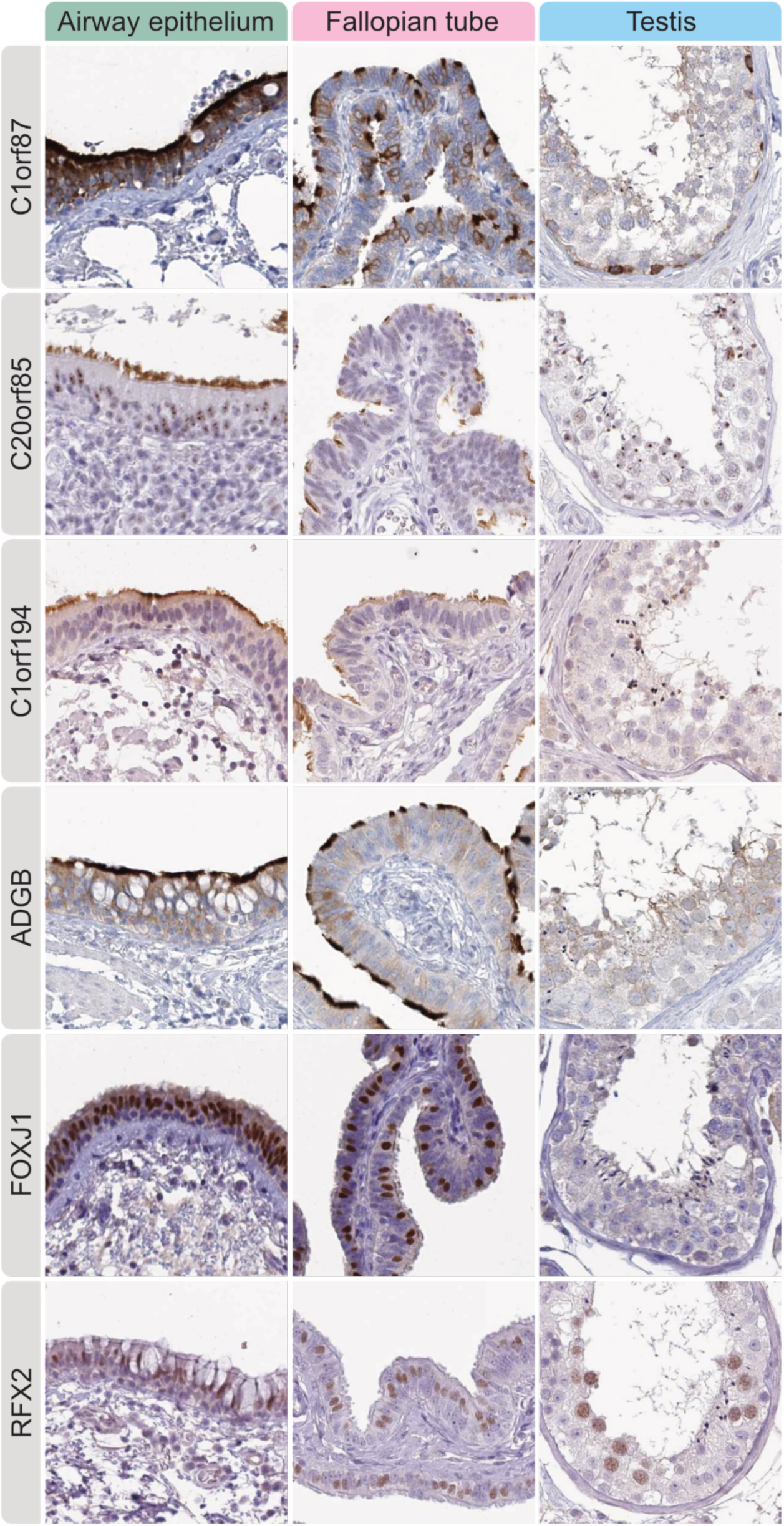
Annotation of the ciliated cell types. Examples of cilia-related proteins differentially expressed in various structures of cilia and/or flagella, with C1orf87 and C20orf85 being expressed both intracellularly in ciliated cells, as well as on the tip of cilia and in ciliary rootlets, both in bronchus and fallopian tube. In testis, C1orf87 stained spermatogonia, while C20orf85 showed dotlike positivity in spermatids and acrosomes. C4orf47 and C1orf194 both displayed distinct immunoreactivity in the tip of cilia, as well as in sperm flagella. ADGB was expressed in cilia axoneme and sperm flagella. FOXJ1 and RFX2 both displayed nuclear staining in ciliated cells of bronchus and fallopian tube, but FOXJ1 lacked expression in testis while RFX2 showed distinct nuclear immunoreactivity in spermatocytes.

**Figure S18.**
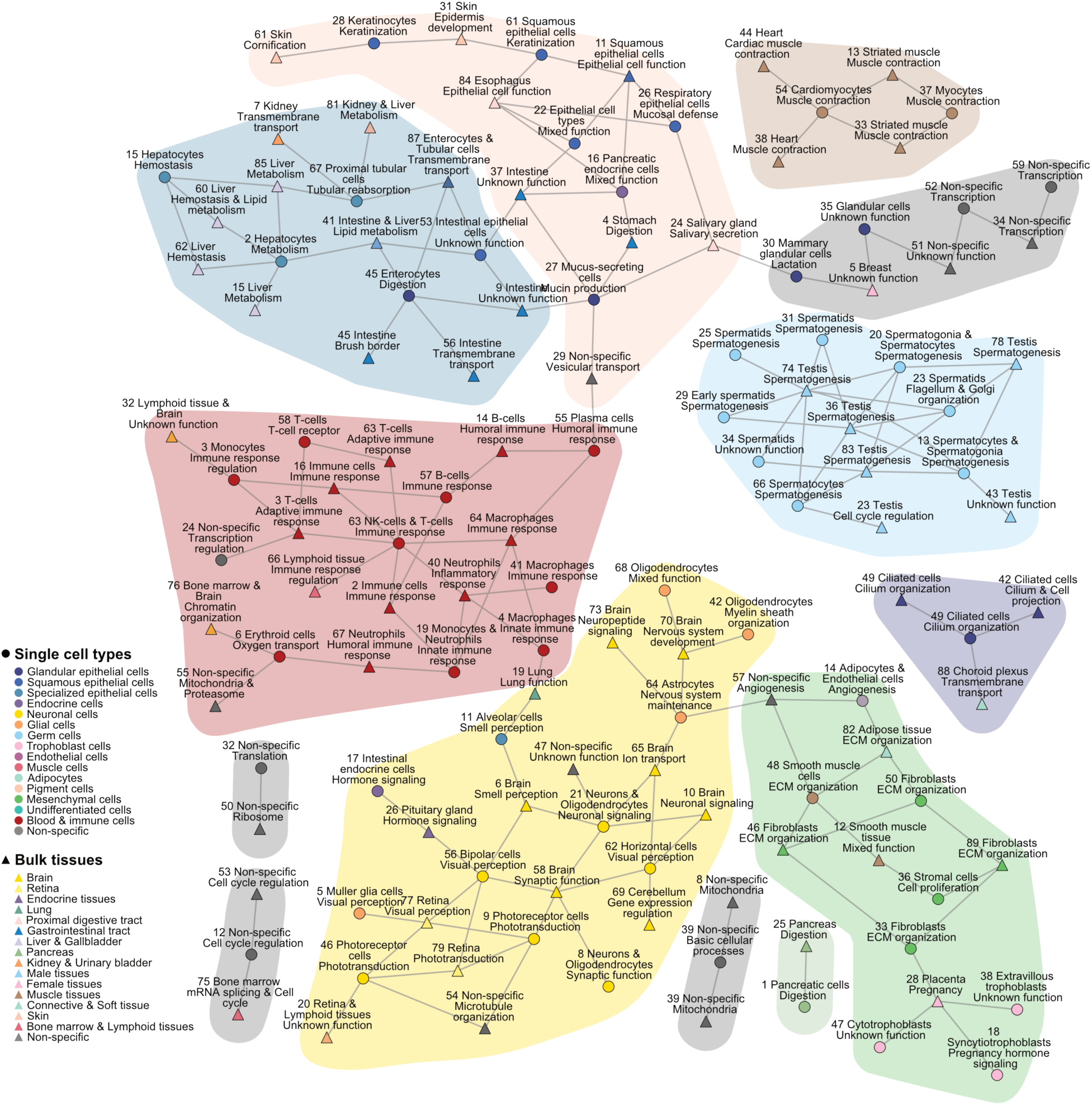
Overlap between genes in the clusters generated from single cell data and bulk tissue data respectively. Links between clusters represent statistically significant gene overlaps (odds ratio > 5, BH adj. p < 0.001), meaning that the connected clusters generated from single cell and bulk tissue data, respectively, contain many of the same genes. Colored areas highlight well connected regions and are colored by the predominant pattern of specificity exhibited by the gene clusters.

**Figure S19.**
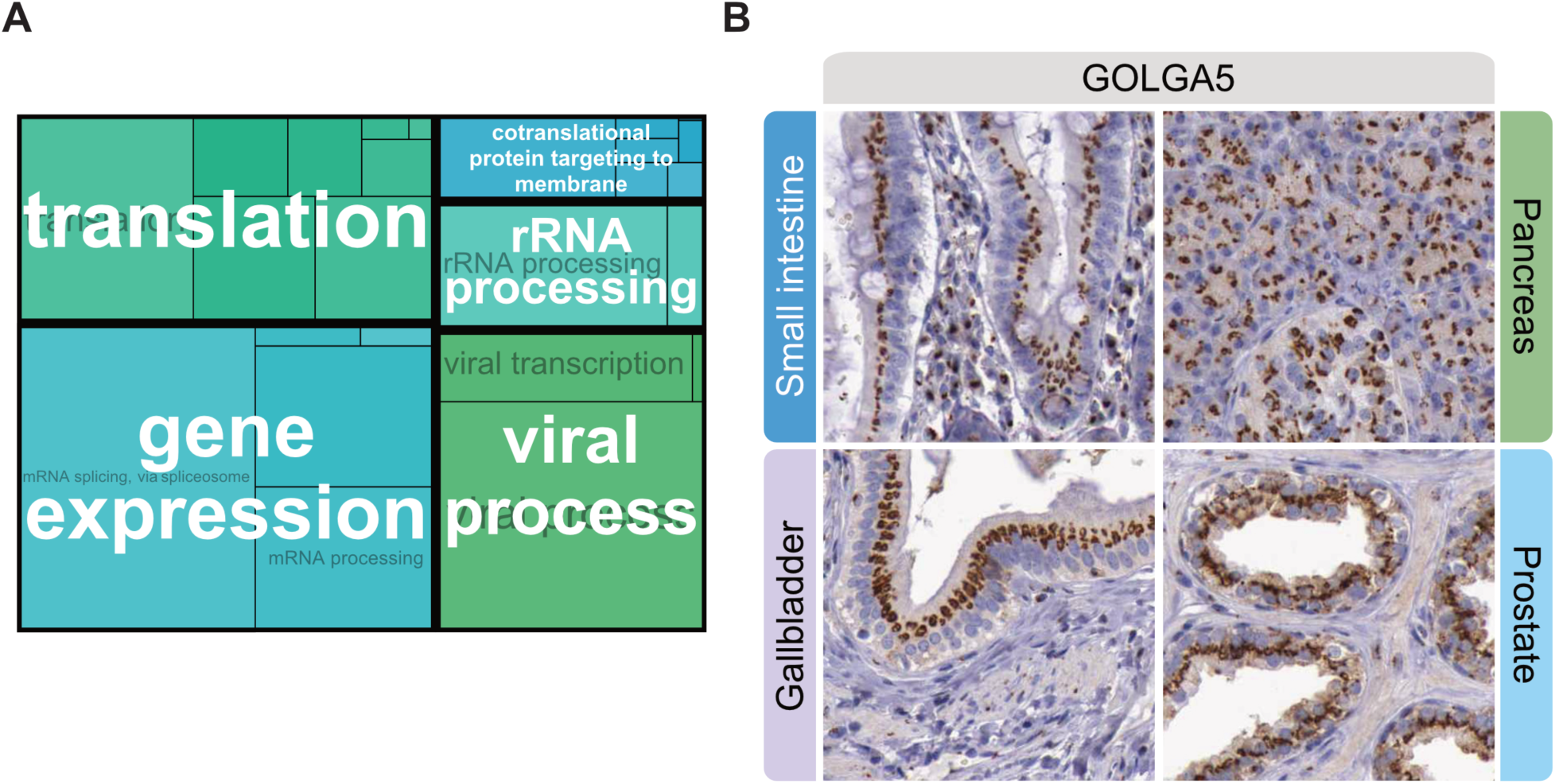
The housekeeping proteome. **(A)** Treemap of GO Biological Process gene set enrichment analysis of the 3230 genes reported as housekeeping in the single cell, immune cells, and cell lines HPA datasets. The GO term box size is proportional to the log-transformed BH adjusted p-values for significantly enriched terms (adj. p < 0.01), and terms with substantial gene overlap are grouped to parent terms. **(B)** IHC image of GOLGA5 showing Golgi-like expression in different tissues.

**Figure S20.**
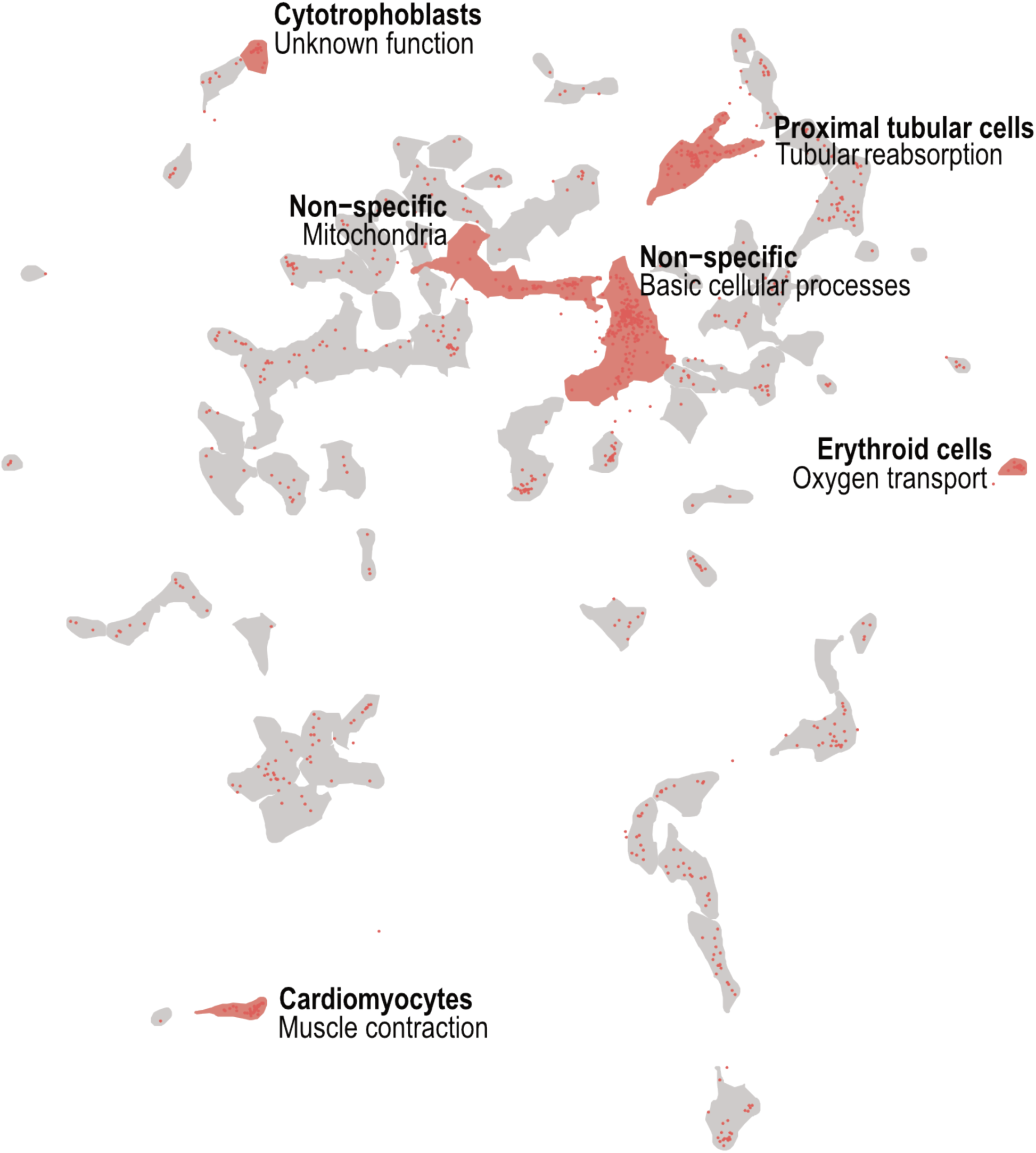
UMAP visualization of protein-coding genes in the single cell data, where genes encoding mitochondrial proteins are highlighted in red (n=947). Clusters with statistically significant overrepresentation (BH adj. p < 0.05) of mitochondrial genes are highlighted in red.

**Figure S21.**
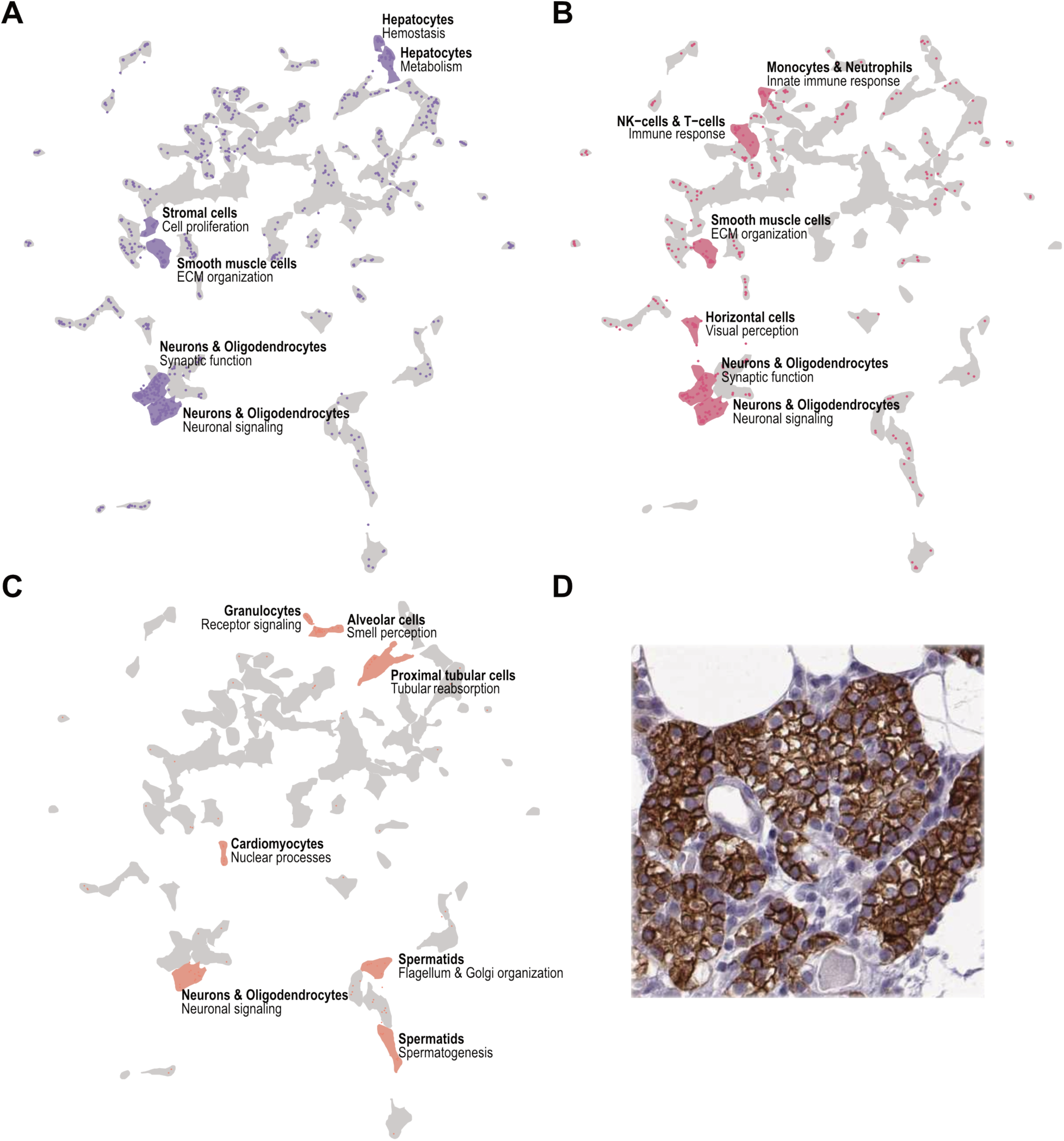
Overview of the druggable proteome. UMAP visualization of protein-coding genes in the single cell data, where genes belonging to the protein classes: **(A)** drug targets, **(B)** non-olfactory GPCRs and **(C)** olfactory GPCRs, respectively, are highlighted in the UMAP. Genes, as well as the clusters with statistically significant overrepresentation (BH adj. p < 0.05) of respective category are highlighted. **(D)** IHC image of the drug target CASR in parathyroid gland.

**Figure S22.**
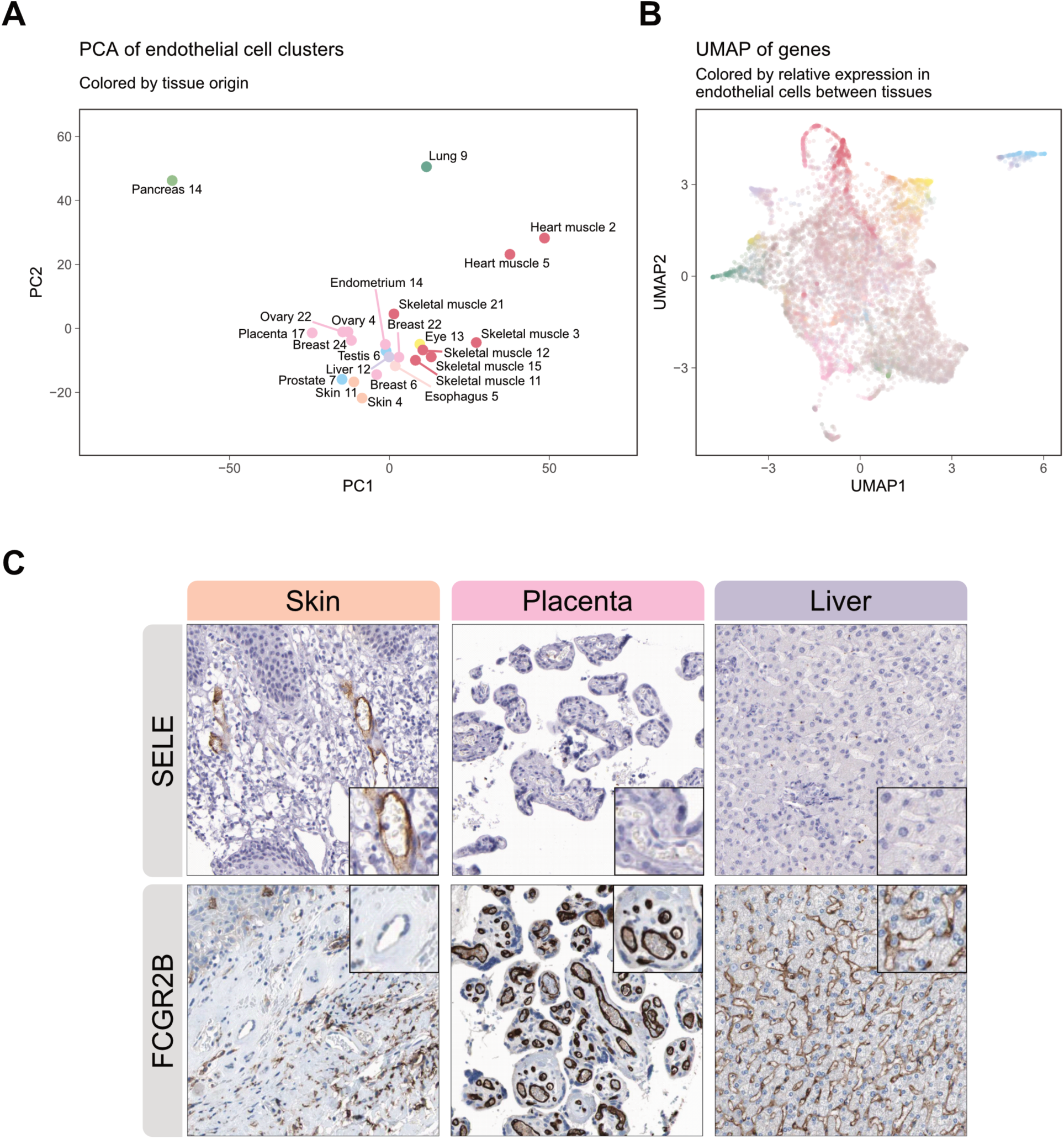
Endothelial single cell type clusters. **(A)** PCA visualization of 23 endothelial cell type clusters (excluding the esophagus cluster) from 13 tissue types. **(B)** UMAP visualization of 16,007 genes (nTPM > 1 in at least 1 endothelial cluster, excluding genes in X and Y chromosomes) based on their expression on the 23 endothelial cell type clusters. **(C)** IHC images of SELE and FCGR2B in skin, placenta, and liver.

**Table S1.** Description of specificity categories in the HPA. nTPM = normalized transcripts per million. Colored circles represent the color associated with each category.

| Category | Description |
| --- | --- |
| ● **Cell type enriched** | nTPM in a particular tissue/cell type at least four times any other tissue/cell type |
| ● **Group enriched** | nTPM in a group (of 2-5 tissues, or 2-10 cell types) at least four times any other tissue/cell type |
| ● **Cell type enhanced** | nTPM in one or several (1-5 tissues, or 1-10 cell types) at least four times the mean of other tissues/cell types |
| ● **Low cell type specificity** | nTPM $\geq 1$ in at least one tissue/cell type but not elevated in any tissue/cell type |
| ● **Not detected** | nTPM $< 1$ in all tissues/cell types |

## Captions for data S1-S6

**Data S1.** List of single cell RNA sequencing datasets

**Data S2.** List of cell type markers used in single cell type cluster annotation

**Data S3.** List of cell types and tissues included in the study

**Data S4.** Cell type classification results of all protein coding genes in terms of specificity and clustering.

**Data S5.** Deep annotation results of cilia related genes.

**Data S6.** Antibodies used for immunohistochemistry

